# Structure of the mature Rous sarcoma virus lattice reveals a role for IP6 in the formation of the capsid hexamer

**DOI:** 10.1101/2020.12.03.410175

**Authors:** Martin Obr, Clifton L. Ricana, Nadia Nikulin, Jon-Philip R. Feathers, Marco Klanschnig, Andreas Thader, Marc C. Johnson, Volker M. Vogt, Florian K.M. Schur, Robert A. Dick

**Affiliations:** Institute of Science and Technology Austria, Klosterneuburg, Austria; Department of Molecular Microbiology and Immunology, University of Missouri, Columbia, MO; Department of Molecular Biology and Genetics, Cornell University, Ithaca, NY

## Abstract

Inositol hexakisphosphate (IP6) is an assembly cofactor for HIV-1. We report here that IP6 is also used for assembly of Rous sarcoma virus (RSV), a retrovirus from a different genus. IP6 was ∼100-fold more potent at promoting RSV mature CA assembly than observed for HIV-1 and removal of IP6 *in vivo* reduced infectivity by 100-fold. By cryo-electron tomography and subtomogram averaging, mature virus-like particles (VLPs) showed an IP6-like density in the CA hexamer, coordinated by rings of six lysines and six arginines. Phosphate and IP6 had opposing effects on CA *in vitro* assembly, inducing formation of T=1 icosahedrons and tubes, respectively, implying that phosphate promotes pentamer and IP6 hexamer formation. Subtomogram averaging and classification optimized for analysis of pleomorphic retrovirus particles revealed that the heterogeneity of mature RSV CA polyhedrons results from an unexpected, intrinsic CA hexamer flexibility. In contrast, the CA pentamer forms rigid units organizing the local architecture. These different features of hexamers and pentamers determine the structural mechanism to form CA polyhedrons of variable shape in mature RSV particles.

## Introduction

Late in the retrovirus lifecycle, the structural protein Gag assembles into an immature lattice at the inner leaflet of the cellular plasma membrane. As the virus buds from the cell, the viral protease cleaves Gag into several fragments, including the capsid (CA) protein that then assembles into the mature capsid shell (core), rendering the virus particle infectious. Experiments based on X-ray crystallography, nuclear magnetic resonance (NMR) spectroscopy, and helical reconstruction have provided information about the structure of CA in isolation and in highly regular tubular arrays [1–5]. In stark contrast, authentic retrovirus particles are pleiomorphic, lacking size and shape uniformity, rendering these techniques not applicable. Cryo-electron tomography (cryo-ET) with subtomogram averaging has proven highly effective in addressing the structure of pleomorphic immature and mature retroviral lattices, optimally resulting in ∼4Å structures [6–9]. An emerging generalization from all of these experiments is that both the immature Gag lattice and the mature CA lattice are based on hexamers of the protein subunits, which exploit different CA interfaces [10]. In most retroviruses, the mature hexameric lattice is closed by incorporation of exactly 12 pentamers, and complete core closure is generally thought to be important for infectivity of retrovirus particles [11]. However, not all retroviral cores form completely closed shells [12]. The CA portion of all retroviral Gag proteins folds into two separate domains, the N-terminal domain (CA_NTD_) and C-terminal domain (CA_CTD_). In mature lattices, CA_NTD_ interactions form only intra-hexamer and intra-pentamer interfaces, while CA_CTD_ interactions are also involved in formation of inter-hexamer and inter-pentamer interfaces.

For viruses in the *Lentivirus* genus, HIV-1 and SIV are highly sensitive to IP6 for immature viral lattice formation and virus release *in vivo* and *in vitro* respectively, and HIV-2, FIV, BIV, and EIAV (immunodeficiency viruses from human, simian, feline, bovine, and equine species, respectively) are sensitive to IP6 for the formation of the immature Gag lattice *in vitro* [13–15]. IP6 is bound to and thereby stabilizes the Gag lattice by interacting with two rings of six lysine residues at the CA-SP hexamer interface. For HIV-1, the mature CA lattice also is influenced by IP6, which binds to a ring of six arginine residues in the N-terminal domain of the CA hexamer. *In vivo*, IP6 is known to be critically important for HIV-1 and SIV replication, since genetic manipulation of cells to remove IP6 dramatically reduces the release of infectious virus particles [13]. *In vitro* assembly of Gag and CA protein into virus-like particles (VLPs) has facilitated study of the lattice structure of many retroviruses [16]. In this methodology, the viral proteins are purified after expression in *E. coli*, incubated under defined conditions of pH, ionic strength, temperature, and in some cases with an oligonucleotide or other additions, and then observed by negative stain electron microscopy. The structures of these VLPs have been shown to accurately mimic those of *bona fide* virus particles [7,17].

Using *in vitro* assembled particles, we here describe the role of IP6 in the mature structure of Rous sarcoma virus (RSV), an alpharetrovirus that has been used widely as a model system for HIV-1 and for studying aspects of cancer biology. Our results show that, like lentiviruses, RSV is sensitive to IP6 *in vitro* and *in vivo*, where in the mature CA lattice IP6 binds in a pore formed in the center of the CA_NTD_ hexamer. Moreover, our subtomogram averaging analysis of pleomorphic polyhedral CASPNC VLPs reveals that a remarkable, previously unobserved flexibility of mature CA hexamers accommodates the wide range of curvatures intrinsic to mature RSV particles and that CA pentamers represent rigid building blocks determining local lattice geometry.

## Results

### IP6 is an RSV assembly cofactor

Like all Gag proteins, RSV Gag comprises multiple domains (Fig. 1a), but as shown by the well-studied truncated protein called GagΔMBDΔPR, only the p10, CA, SP, and NC domains are required for efficient immature virus particle assembly *in vitro* [18,19]. In immature assembly in lentiviruses, conserved lysine residues in the major homology region (MHR) and CA-SP1 helix bind IP6 [15] (Fig 1a). These residues, however, are not present at the corresponding positions in RSV Gag. Consistent with this lack of sequence conservation, RSV GagΔMBDΔPR supports efficient *in vitro* assembly of immature VLPs in the absence of IP6 (Fig. 1b). At a physiological 50 µM concentration of IP6, assembly was not affected. However, at higher concentrations, IP6 led to decreased or abrogated assembly. As described below, this inhibition of immature assembly may be due to the strong effect of IP6 on mature assembly.

**Fig. 1:**
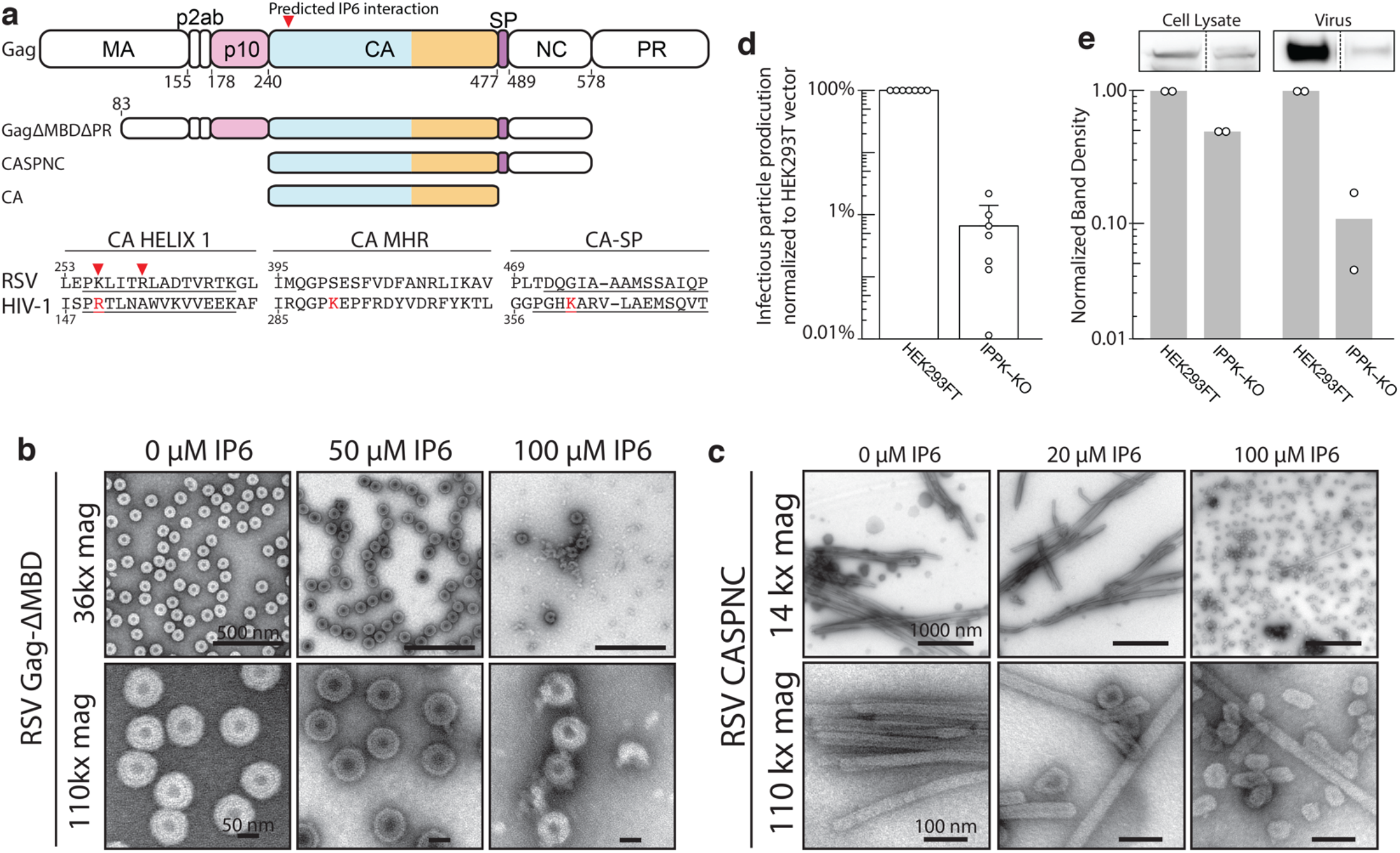
Effect of IP6 on RSV in vitro and in vivo. **a)** RSV Gag and truncation proteins used for assembly and structure determination. Alignment of RSV and HIV-1 amino acid sequences. Known HIV-1 IP6 interacting residues shown in red and predicted RSV IP6 interacting residues indicated (red arrow). **b)** TEM images of immature RSV Gag-ΔMBD assembly without and with IP6 at low and high magnification. **c)** TEM images of mature RSV CASPNC assembly without and with IP6 at low and high magnification. **d)** RSV infectious virus particle production in wild type (HEK293T) and IPPK-KO cell lines. **e)** Representative western blots (top) and the average of two western blot experiments of cell lysate (left) and virus fractions (right). Western blots are from the same gel, cropping indicated by box and dotted lines.

RSV CASPNC, a construct similar to GagΔMBDΔPR, but missing the p10 domain that is known to be critical for immature lattice formation, assembled into mature long narrow tubes, consistent with previously published results [20]. The addition of IP6 to this protein resulted in a dramatic change in morphology, from tubes to (non-uniform) polyhedrons that resemble mature cores in the infectious virus particles [21] (Fig. 1c). These polyhedral VLPs appeared first in significant numbers at 20 µM IP6, and then dominated the population of VLPs as the IP6 concentration was raised to 100 µM. Consistent with this effect of IP6 on mature assembly, RSV CA has lysine and arginine residues in Helix 1 of CA, located near the site of the HIV-1 R18 residue that interacts with IP6 at the mature CA_NTD_ hexamer pore (Fig. 1a).

Previously we showed that depletion of IP6 from HEK293FT cells by CRISPR-Cas9 knock-out of inositol-pentakisphosphate 2-kinase (IPPK), the enzyme that converts IP5 to IP6, caused a ∼20-fold reduction in the production of infectious HIV-1 particles. In the case of RSV, infectious RSV production was reduced ∼100-fold from HEK293FT IPPK-KO compared to WT without the need for further genetic manipulation (Fig. 1d). These results show that IP6 plays an important role for infectious particle production of RSV, but do not clarify at what step or steps IP6 acts.

To delineate how the removal of IP6 impacts RSV, we performed western blot analyses of whole cell lysates as well as the supernatants to measure relative levels of viral protein produced and virus particles released (Fig. 1e). Quantification showed that Gag synthesis was reduced nearly 2-fold in HEK293T IPPK-KO cells compared with WT HEK293T cells (Fig 1e, left). Virus release was reduced by ∼10-fold in IPPK-KO cells (Fig. 1e, right). We interpret these results to mean that while RSV is at least partially dependent on IP6 for Gag translation in cells, the major effect of IP6 removal is to reduce the assembly and release of virus particles from cells. These results are the first to indicate IP6 as an assembly co-factor of a retrovirus outside of the *Lentivirus* genus.

### Structure determination of the mature RSV lattice

We sought to structurally characterize the mature RSV CA lattice to better understand how IP6 influences mature assembly. We acquired cryo-ET data of CASPNC VLPs assembled in the presence of 100 µM IP6. The reconstructed tomograms of this sample showed abundant regular tubes and pleomorphic polyhedral VLPs. (Fig. 2a, Movie S1). Conventional subtomogram averaging, as described previously [14], of tubular VLPs with regular diameter and shape resulted in a reconstruction of the CA hexamer at 4.3 Å resolution (Fig. 2b, left, Fig. S1a, Fig. S2a). The EM density map revealed a similar arrangement of the CA_NTD_ and CA_CTD_ as previously reported for other mature retroviral VLPs (Movie S2). Given the inherent two-fold symmetry of the tubes, CA monomers within tubes are present in three symmetry-independent copies. In the center of the CA_NTD_ hexamer an additional density is present in the identical position, which has previously been reported as an IP6 binding site in mature HIV-1 [15]. No ordered density for NC and also no ordered density for the beta-hairpin, which has been observed in other retroviral mature CAs, were observed. At the resolution of 4.3 Å, we were able to refine a model of RSV CA [22] into the EM density of the mature hexameric CA assembly. As reported for other mature retrovirus VLPs, intra-hexameric interactions are established via CA_NTD_-CA_NTD_ as well as CA_NTD_-CA_CTD_ interfaces, while the inter-hexameric interactions are formed solely via CA_CTD_-CA_CTD_ interactions maintaining contacts across two-fold and three-fold interfaces [1,3,9,22]. Notably, an IP6 molecule could be fitted into the density in the CA_NTD_ hexamer center in an upright position, where it is coordinated by two concentric rings of positively charged residues K17 and R21 (Fig. 2d, Movie S2).

**Fig 2.**
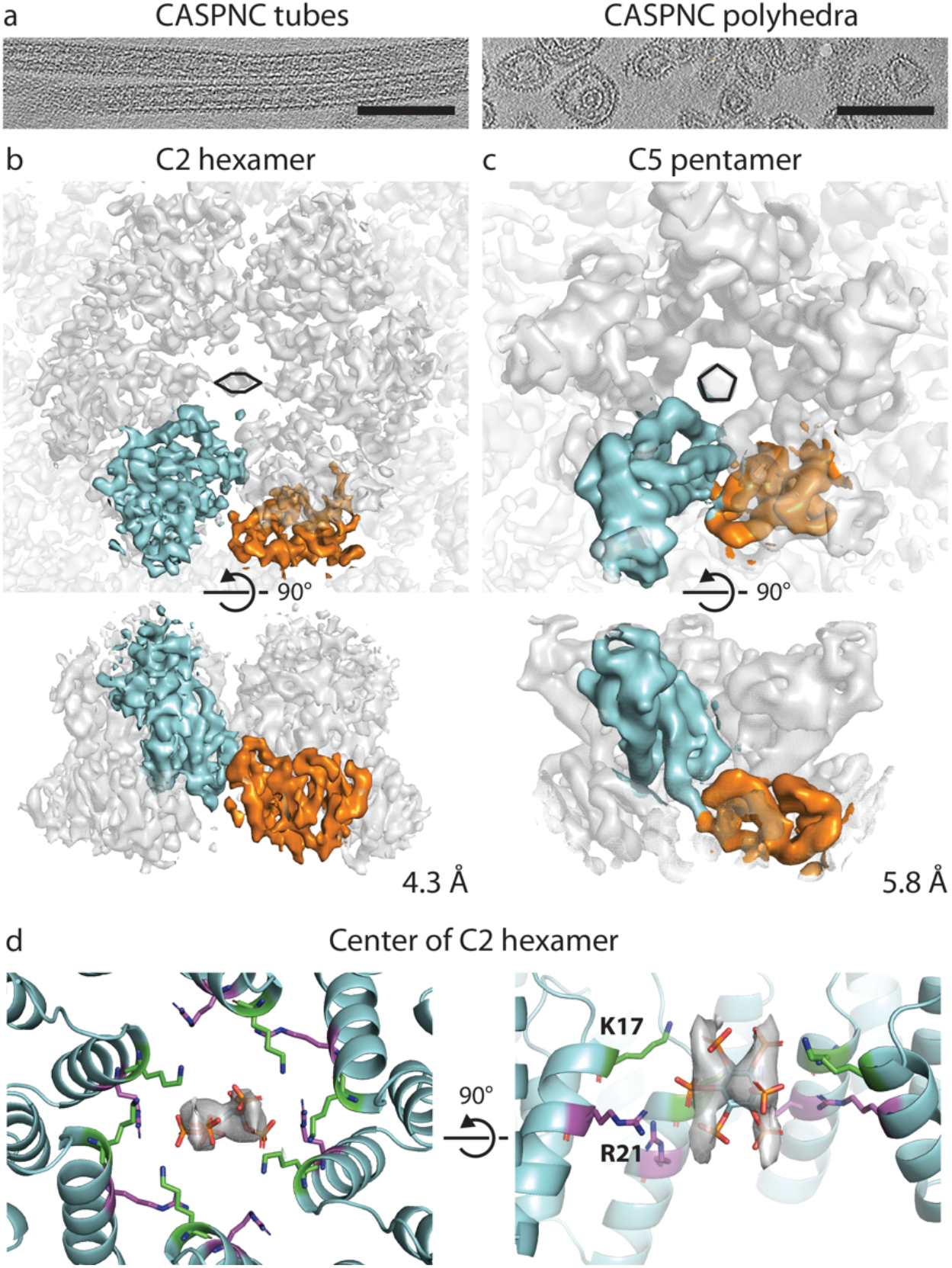
Cryo-electron tomography of RSV CASPNC VLPs assembled in the presence of IP6. a) Sum of 10 computational slices through a gaussian-filtered tomogram containing CASPNC tubes and CASPNC polyhedrons. Scale bar is 100 nm. **b-c)** Isosurface representation of the subtomogram average of a C2-symmetric CA hexamer from the CASPNC tubes (b) and a C5-symmetric pentamer from CASPNC polyhedrons (c). In both cases, the CA_NTD_ and CA_CTD_ of one CA monomer are colored cyan and orange, respectively. The resolution is annotated. The C2 and C5 symmetry of the hexamer and pentamer are annotated by a distorted schematic hexamer and a pentamer, respectively. **d)** Model of the IP6 binding site in the CA-NTD hexamer pore formed by amino acids K17 (green) and R21 (purple) as seen from the outside of the VLP (left) and rotated by 90° (right).

### Structural determinants of pleomorphic mature RSV CA lattices

To further understand how RSV CA is able to generate both regular tubular and pleomorphic polyhedral VLPs, and in particular how the different inter- and intra-hexameric CA interfaces allow for the observed flexibility, we also performed subtomogram averaging on polyhedral VLPs. Most important, we determined the structure of the mature CA pentamer in these polyhedrons to define the interactions between hexameric and pentameric RSV CA assembly states.

The highly variable size and shape of RSV CA polyhedral VLPs, ranging from small icosahedrons and polyhedrons to several micrometer long tubes (Fig. 2a), resulted in a variable and limited local order of the CA lattice. A conventional subtomogram averaging approach (Fig. S1a, Fig. S4a) assumes a high degree of rigidity and symmetry of the CA oligomeric units (i.e. hexamers and pentamers) and an extended local order of the CA lattice, which implies that the variations in the local curvature come primarily from varying interfaces between CA hexamers or pentamers. However, as conventional subtomogram averaging of polyhedral RSV CASPNC VLPs only yielded a structure of the CA hexamer with limited resolution, despite a significant number of asymmetric units, we concluded that these described geometric conditions were not met in RSV CA polyhedral VLPs and that hexamers and their surrounding show significant flexibility and variability.

Hence, in order to appropriately address the plasticity of the polyhedron CASPNC lattice and any potential conformational flexibility within individual CA hexamers, which could be associated with their different local environments (such as a varying curvature), we devised a novel subtomogram averaging approach that uses classification and alignment guided by local geometry and contextual information (Fig. S1, Fig. S3, see also *Materials and Methods*). In order to classify hexamers and pentamers according to their geometrical context, we defined two criteria that we found to influence the geometry of the mature CA lattice: 1) the presence and location of one or more pentamers in the vicinity of a hexamer or pentamer (Fig. S3a), and 2) the orientation of a pair of just hexamers, a pair of hexamers and pentamers or a pair of just pentamers (termed here as unit pairs) with respect to the local curvature (Fig S3b). According to these criteria all hexamers and pentamers were classified into different classes belonging to 6 general groups (Fig. S3c).

For determining the structure of the asymmetric parts of lattice we implemented an alignment strategy respecting the asymmetry of each unit given by its local environment. Specifically, instead of treating hexamers and pentamers within the RSV CA lattice as isolated symmetric units and using a single alignment against a common reference, the alignment of each hexamer or pentamer in the mature lattice is decomposed into several independent alignments (Fig. 3a and b, Fig. S4b, see *Materials and Methods* for further details). In brief, each hexamer and pentamer is aligned multiple times (i.e. 6 times in case for a hexamer that is having six neighbors), each time using a reference specific to its local context, and hence geometry. The consensus of these separate alignments then determines the position and orientation of the respective oligomeric unit (Fig. S4b). In the final step, an additional alignment centered in the middle of the unit pair (i.e. the dimer interface) was performed, in order to optimize the structures of different classes with respect to the interfaces between the unit pairs (Fig. S1b).

**Fig 3.**
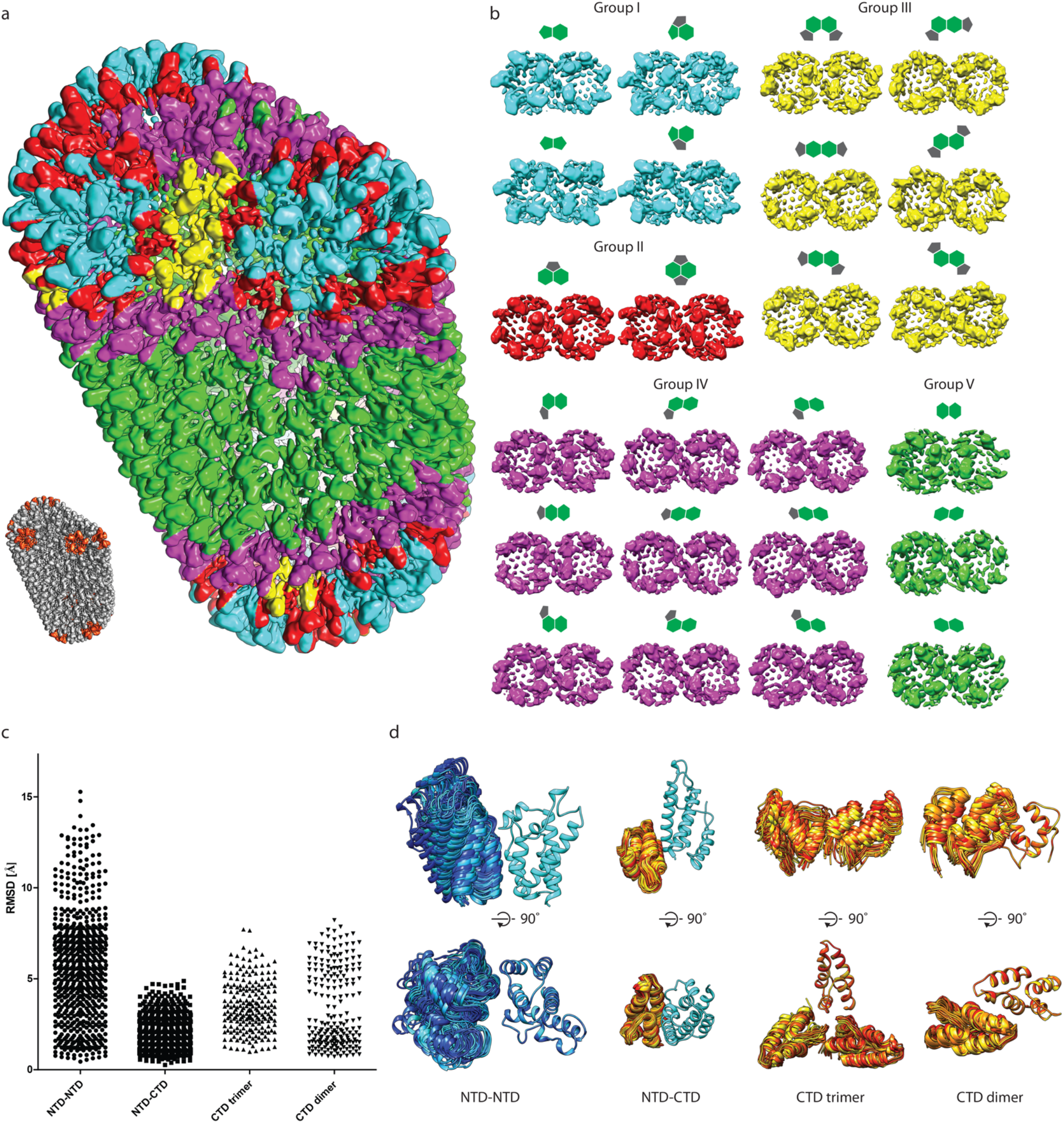
Building blocks of the mature CASPNC lattice. **a)** A composite model of a representative RSV CASPNC VLP, generated by placing respective unit pair averages (see groups in b) into their positions derived from subtomogram alignment. **b)** Final averages of unit pairs filtered to 8 Å. Groups are identical to their schematic depiction in Figure S3. Coloring of the groups corresponds with panel a. **c)** Pairwise RMSD measurements between rigid-body fitted models derived from the maps of the different classes shown in panel (b). In case of CA_NTD_-CA_NTD_, CA_NTD_-CA_CTD_ and CA_CTD_-dimer interfaces, one CA domain is fixed and RMSD is measured for the other CA domain across the respective interface. For the CA_CTD_-trimer interface one CA domain is fixed and RMSD is measured for the other two CA_CTD_’s forming the trimer interface. **d)** Superimposition of the models showing the flexibility of CA domains involved in the different interfaces. Color code: cyan to blue – CA_NTD_; yellow to red – CA_CTD_.

This approach yielded 21 distinct structures at resolutions of 6-9.1Å of RSV hexamer/pentamer, hexamer/hexamer and pentamer/pentamer pairs that exist in mature RSV CA lattice (Fig 3b, Fig. S2b) and also yielded a structure of the CA pentamer at 5.8 Å (Fig. 2c, Fig. S2a). For further analysis we rigid body-fitted individual CA domains into the EM-density maps of hexamer/pentamer unit pairs. These structures and models indeed confirmed that the mature polyhedron CA lattice consists of structurally different, intrinsically flexible CA hexamers that strongly deviate from regular six-fold symmetry. In other words, the hexamers need to accommodate varying interactions and curvatures to form the observed highly pleomorphic polyhedrons (Fig. 3a and b, Movie S3). Hexamers which are separated from pentamers by two or more hexamers, are essentially identical to those found in regular tubes. Pentamers, on the other hand, organize rigid vertices, where the lattice is curved in different directions. The hexamers that are adjacent to pentamers are deformed to allow bridging of the tube-like areas and the pentamer vertices. (Fig. 3a and b).

Further analysis of the different hexamers forming the mature CA lattice in polyhedral VLPs revealed that despite the deformability of the CA hexamer, local symmetries exist in the lattice (Fig. S2a). Aside from the C2-symmetric hexamers that are observed in regular tubes (and are surrounded by only hexamers), a fraction of hexamers in the polyhedrons were surrounded by either two or three pentamers in a symmetrical manner and hence follows C2-symmetry and C3-symmetry, respectively (Fig. S5 a and b). As the structure of the tube-like hexamer with C2 symmetry was already solved at high resolution from regular tubes, it was not further characterized. In contrast, the structures of the other two symmetric hexamers highlight the structurally dominant influence of the pentamer, where it actively defines the conformation of its adjacent hexamers.

Specifically, the two CA monomers facing a pentamer adopt an arrangement similar to those two CA monomers in the pentamer, causing an increased spatial separation between the CA_NTD_’s of the hexamer (Fig. S5 a to d, Movie S4). This observation was further confirmed via the structure determination of a small set of T=3 icosahedral particles solved by single particle cryo-EM (Fig S1c, Fig. S5e). In T=3 icosahedrons each hexamer is surrounded by three pentamers, which causes the described three-fold symmetric distortion of the CA hexamer. The similarity of the C3 hexamer from the T=3 icosahedra (formed by CA) to the C3 hexamer from the polyhedrons (formed by CASPNC and IP6) (Fig. S5 f and g) suggests that the structural deviation from C6 symmetry is not due to the presence of either the IP6 molecule, nor the absence of the β-hairpin, nor the SPNC region.

The finding of the strong plasticity of the mature RSV CA hexamer led us to compare the variability of the intra-hexamer and intra-pentamer CA_NTD_-CA_NTD_ and CA_NTD_-CA_CTD_ interfaces, as well as the dimeric and trimeric CA_CTD_ interfaces in our structures using pairwise RMSD analysis. (Fig 3c and d). In agreement with the observed deformability of the CA_NTD_ ring in hexamers, CA_NTD_-CA_NTD_ interfaces displayed the largest variation. In contrast, CA_NTD_-CA_CTD_ interactions are locking the two domains in a quaternary arrangement that does not deviate extensively between the different classes. Two populations of CA_CTD_ dimer interfaces (termed here as state A and B) were observed (Fig. 3d, Fig. S6, Movie S5). State A was found in all dimer interfaces involving pentamers (with higher curvature) and low curvature hexamer-hexamer interfaces, whereas state B was found only in hexamer-hexamer interfaces with high curvature. Specifically, comparison of high-resolution structures of T=1 particles (see below) and one hexamer-hexamer interface from regular tubes revealed striking similarity (Fig. S6a, Movie S5). The changes of the dimer interface in the two states had impact also on the CA_CTD_ trimer interface, and induced its deformation from C3 symmetry. (Fig. 3c and d).

This interface analysis explains how the rigid CA_CTD_-CA_NTD_ interface acts as a structural link to propagate the changes in the CA_CTD_ dimer interface towards the hexameric CA_NTD_ ring to distort the weak interactions across the CA_NTD_-CA_NTD_ interface.

### IP6 shifts phosphate-induced CA VLP assembly from pentameric to hexameric

The addition of IP6 to CASPNC assembly reactions caused a dramatic shift from tubular to polyhedral VLPs. Since polyhedrons contain both hexamers and pentamers while tubes contain only hexamers, a possible interpretation of this result is that IP6 promotes pentamer formation. An alternative interpretation is that IP6 promotes initiation of assembly, which ultimately results in rapid depletion of CA. This rapid depletion of CA might favor polyhedron assembly. To shed light on these possibilities, we performed *in vitro* assembly of CA protein with IP6 and/or NaPO_4_. High concentrations of NaPO_4_ are known to induce RSV CA to assemble predominantly into T=1 icosahedral VLPs, which consist of only twelve pentamers. If IP6 promotes pentamer formation this small molecule would be expected also to induce assembly of icosahedral VLPs. On the other hand, if IP6 promotes hexamer formation, we predicted that it would promote tubular VLPs. The results of this experiment showed that in absence of IP6 or NaPO_4_, CA appeared in negative stain EM images as small aggregates and unassembled protein (Fig. 4a). But in the presence of 100 µM IP6, CA assembled into multilayered tubes. Raising the level of IP6 to 500 µM led to a transition to single layered tubes (Fig. 4a). Higher concentrations of IP6 resulted in protein aggregation. We interpret this result to mean that IP6 favors the formation of CA hexamers over CA pentamers.

**Fig 4.**
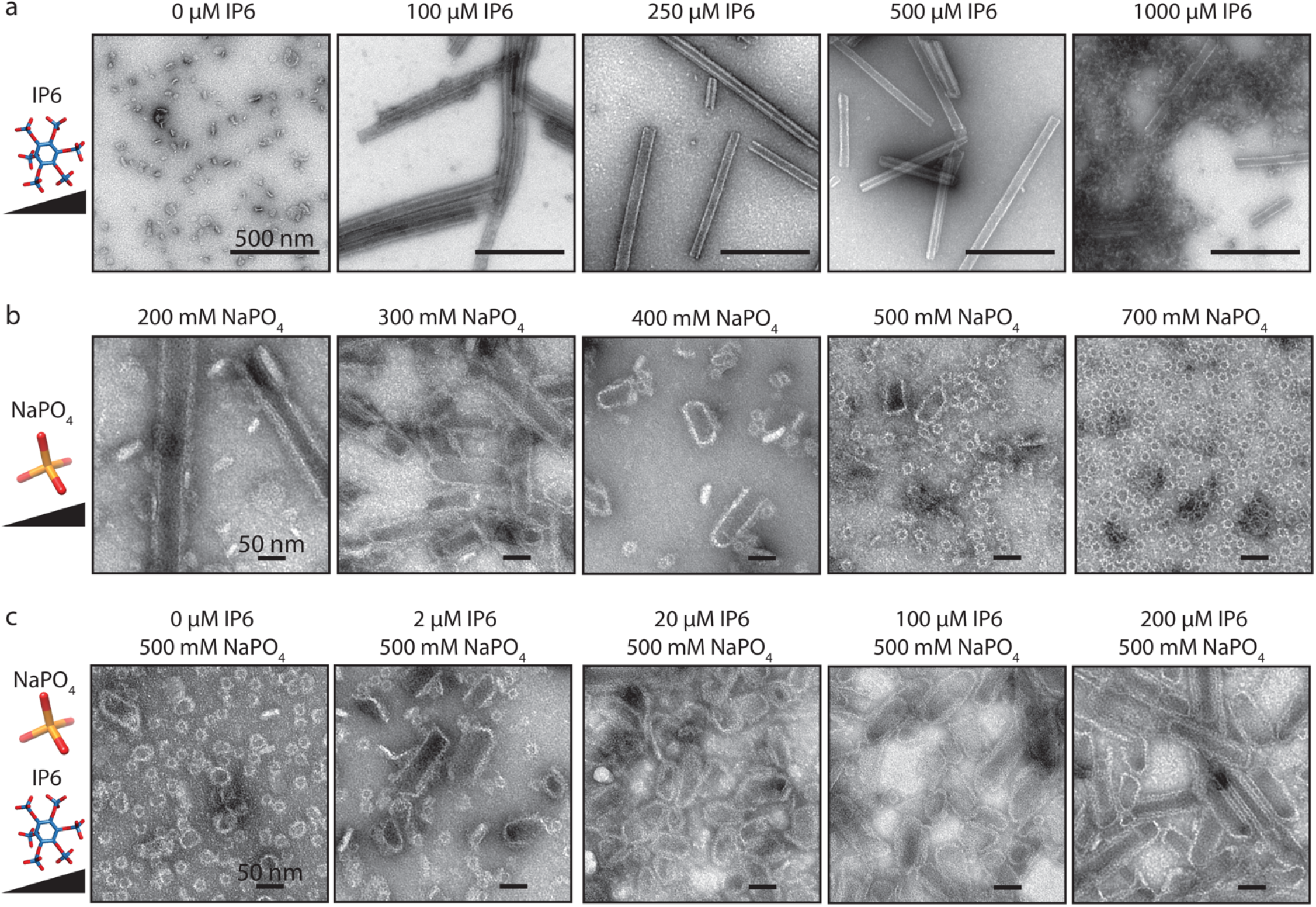
Effect of NaPO_4_ and IP6 on in vitro CA assembly. **a)** Example TEM images of CA assembly reactions with increasing concentrations of IP6. **b)** Example TEM images of CA assembly reactions with increasing concentrations of NaPO_4_. **c)** Example TEM images of CA assembly with fixed 500 µM NaPO_4_ and increasing concentration of IP6.

High phosphate concentrations were previously predicted to affect assembly *in vitro* by shielding the charges of the K17 and R21 of the CA pentamer [23,24], allowing these side chains to come into close proximity in the CA pentamer. In the presence of 200 µM NaPO_4_, CA assembled into some multilayered tubes (CA hexamers), but also considerable aggregates (Fig. 4b). In presence of 300 mM NaPO_4_, CA formed predominantly short tubes with curved ends, which we interpret to reflect the presence of CA pentamers. At 400 mM NaPO_4_, T=1 icosahedral VLPs were observed mixed with blunt-ended tubes. The number of T=1 VLPs increased as NaPO_4_ was increased, and at a concentration of 700 mM nearly all of the particles were small icosahedral VLPs. If phosphate favors pentamer formation and IP6 favors hexamer formation, we speculated that the addition of IP6 to assembly reactions that normally result in icosahedrons could shift the resulting VLPs from predominantly icosahedral to predominantly blunt tubes. While at 500 mM NaPO_4_ CA formed predominantly small icosahedral VLPs, the addition of 2 µM IP6 resulted in an observable shift to fewer icosahedrons and more polyhedrons and blunt-ended tube VLPs (Fig. 4c). At 20 and 100 µM IP6 the majority of VLPs were polyhedrons. At 200 µM IP6 the majority of CA assembled into blunt-ended tubes. Based on these results, we conclude that IP6 promotes assembly by interacting primarily at the CA hexamer interface.

Based on the *in vitro* assembly and structural data, IP6 and NaPO_4_ likely promote assembly via interactions with K17 and/or R21 amino acid side chains. To verify this, CASPNC protein with either K17A or R21A mutations was purified and tested for assembly. Compared to WT (Fig. 1c) CASPNC K17A formed fewer tubes, and many small polyhedral VLPs in the absence of IP6 (Fig S7a). The addition of IP6 decreased the number of polyhedrons, with assembly favoring tubes. CASPNC R21A protein was assembly incompetent both without and with IP6 (Fig S7a). We also tested the effect of the K17A mutation on CA assembly. In the absence of IP6 or NaPO_4_, CA K17A formed some aggregates and thin walled assemblies (Fig S7b). Assembly in the presence of IP6 resulted in the formation of small (∼10 nm) cross-structures, which we interpret to be a regular CA complex that does not support mature lattice formation. Assembly reactions performed in the presence of NaPO_4_ resulted in narrow tubes (∼10 nm in diameter) and the same cross-like structures. Both K17A and R21A mutations abrogated immature assembly of the GagΔMBDΔPR protein (Fig S7c), likely due to the mutation disrupting an interface critical to the immature lattice. Taken together, we interpret these results to mean that the charge of K17 and its neutralization (by IP6 or NaPO_4_) are critical factors regulating RSV assembly.

### CA pentamer structure from the RSV T=1 icosahedron

The observation that within pleomorphic polyhedrons the CA pentamer constitutes the rigid building block prompted us to investigate if the pentamers within T=1 icosahedrons and polyhedrons are structurally identical. Hence, we acquired cryo-EM micrographs of T=1 icosahedral VLPs and subjected them to single particle analysis, yielding a 3.1Å resolution reconstruction (Fig. 5a-c). At this resolution refinement of RSV CA into the EM-density was unambiguously possible (Fig. 5 d). We compared the model of T=1 pentamers with the model of the pentamer derived from the polyhedrons. In fact, the two models are virtually superimposable, showing an RMSD of their C-alpha atoms of 0.74 Å (Fig. 5e). This finding further underscores the observations that the RSV CA pentamer (1) is very rigid due to the tighter packing of the CA domains in the pentamer, (2) always promotes the same local geometry, and (3) is independent in arrangement of the size and shape of the VLP.

**Fig 5.**
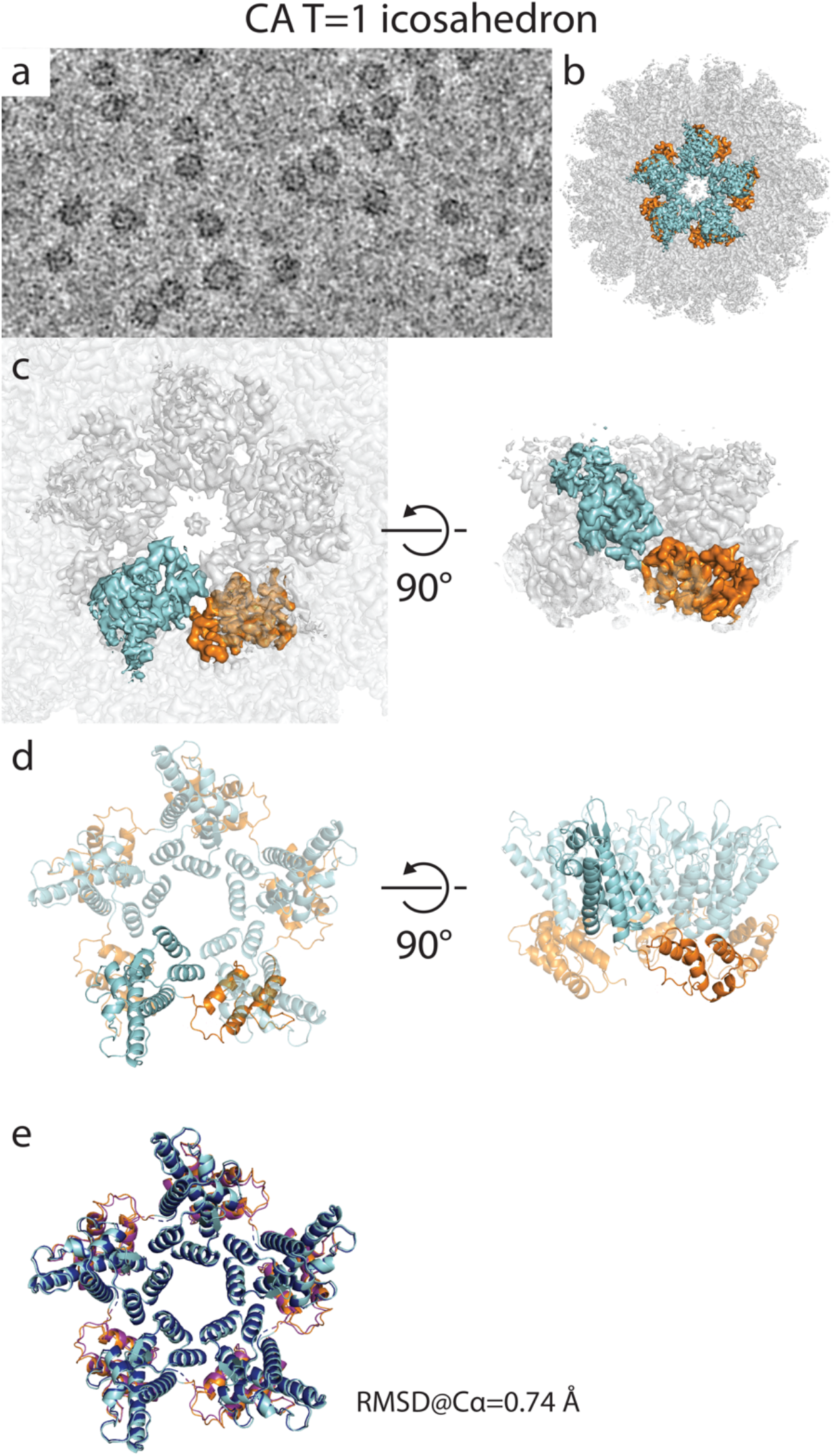
Structure of the RSV T=1 icosahedron pentamer. **a)** Left: filtered cryo-EM micrograph of T=1 icosahedrons. Right: Isosurface representation of the CA T=1 icosahedron determined by single particle cryo-EM. One pentamer is highlighted in colors: cyan – CA_NTD_; orange – CA_CTD_. **b)** Isosurface representations of one T=1 pentamer from the structure shown in (a). **c)** Model of the RSV CA pentamer refined from the structure shown in panel (b). **d)** Comparison of the real-space refined model of the CA pentamer from T=1 particles and rigid-body fitted model of CA pentamer from polyhedrons. The low RMSD value of their backbone atoms shows that the two models are almost identical.

## Discussion

IP6 is an assembly cofactor for HIV-1 and other lentiviruses [13–15,25]. Here we have demonstrated that IP6 has a potent effect on the assembly and morphology of the mature RSV CA lattice, and that depletion of IP6 from cells inhibits virus release and spread. This is the first study to demonstrate that IP6 is an assembly co-factor for a virus not in the *Lentivirus* genus. In addition, we have determined the structure of the mature RSV CA hexamer and pentamer at the highest resolution to date, using cryo-ET and cryo-EM, respectively, to reveal structural mechanisms underlying RSV mature core formation.

The effect of IP6 on mature RSV assembly is unambiguous. The interaction of IP6 with K17 and R21 is similar to the interaction of IP6 with R18 in HIV-1 [15]. For HIV-1, the IP6 molecule sits above the top of the pore at residue R18 (but can also be found below the R18 residue, effectively in the pore), whereas for RSV IP6 is found in the pore interacting with residues K17 and R21. Most retroviruses have an arginine or lysine residue in the first helix of CA, corresponding to K17 and R18 of RSV and HIV-1, respectively. The charged pore of the mature hexamer is thought to be used for dNTP import, with IP6 acting there to overcome the repulsive charge of closely positioned arginine and/or lysine residues during assembly, as proposed by [23,24]. The only orthoretroviruses that do not have basic residues in helix-1 belong to the *Gammaretrovirus* genus, which includes murine leukemia virus (MLV). From a recent cryo-ET structure of the mature MLV lattice, it appears that unlike lentiviruses and alpharetroviruses, gammaretroviruses do not form a closed capsid shell [12], thus possibly allowing dNTP import through gaps in the lattice. If this model is correct, IP6 would be important for the assembly of retroviruses from many different genera.

While IP6 significantly enhances the immature assembly of lentiviruses [13–15], we did not observe a similar effect for RSV. The lentiviral immature IP6 binding site consists of two rings of six lysine residues each. For HIV-1 this is K290 (located in the MHR) and K359 (located at the C-terminal end of CA, in the first turn of the helix that forms the six-helix bundle). RSV does not have basic residues located at or near its MHR or the putative six-helix bundle. Nevertheless, we found that depletion of IP6 from cells reduces virus particle release. Thus, IP6 either directly influences immature assembly via a Gag-IP6 interaction, or indirectly influences assembly or release via some yet to be determined mechanism.

For HIV-1, IP6 binding to the immature interaction site at K290 and K359 in the Gag lattice ensures packaging of IP6 into the budding virus particle. Following maturation, IP6 thus is already present to promote mature assembly via interactions with the mature hexamer at R18. If RSV does not directly require IP6 for immature assembly, how is IP6 incorporated into the virus particle? Sensitivity to IP6 during mature assembly was 100 to 1000-fold greater for RSV than for HIV. Perhaps a small amount of IP6 is taken into the budding RSV particle non-specifically, and this amount is sufficient for the formation of the mature lattice. Alternatively, perhaps in RSV proteolytic maturation of Gag occurs during budding, allowing IP6 to be incorporated by the growing mature CA lattice before the viral membrane has been sealed. The genome structure of RSV is unusual in that the viral protease is part of the Gag polyprotein and possibly allowing PR to become active earlier than for other retroviruses.

Retroviruses from different genera adopt varying quaternary CA_NTD_ arrangements in the immature Gag lattice [10]. In contrast, no significant deviations have so far been reported for quaternary CA arrangements in mature retroviruses. The most notable differences instead were observed on the global architecture of the mature CA lattices, with HIV-1 forming cone-shaped cores with a defined incorporation of pentamers [26] and MLV forming multi-layered, spiral structures with irregular pentamer distribution [12]. In authentic RSV virions mature CA cores form highly pleomorphic structures, either having continuous “spherical” curvature, or tubular and fullerene-like cones shapes [21]. This remarkable core polymorphism is recapitulated by our mature RSV CASPNC and CA *in vitro* assembly, which exhibits a range of differently sized VLPs. The exact mechanism of how mature retroviral CA assemblies are able to form architectures with such a large variety of different curvatures and molecular interactions, is not yet entirely understood. Recent studies analyzing mature HIV-1 CA tubes suggested that the plasticity of the CA hexamer, specifically its deviation from 6-fold symmetry, defines curvature adaptation in mature cores, without significantly altering the CA_CTD_ dimer and trimer interfaces [4,5].

Our results, obtained via a novel classification and alignment approach accounting for asymmetry of CA units in polyhedron assemblies, now extend these findings and provide a more detailed understanding of the dynamic CA properties that underlie curved retroviral CA lattices.

In line with the observations made in HIV-1 [4], mature RSV CA forms pseudo-symmetric, intrinsically curved hexamers. However, due to the highly pleomorphic shape of our VLPs, the structural plasticity of the RSV hexamer is significantly more pronounced, resulting in a large structural variety of hexamers adopting 2-fold and 3-fold symmetric assemblies, as well as asymmetric structures, which are distorted by local curvature and geometrical context. Hence, while hexamers are building blocks of any mature CA lattice, their intrinsic structure and flexibility needs to vary to adapt different requirements in mature core formation. The observed increased flexibility of the intra-hexameric contacts corresponds with increased rigidity of inter-unit interfaces. However, the degree of curvature variability in the mature lattice still requires different connections between the units. Hence, instead of having a continuum of different inter-hexameric and pentamer interactions gradually adjusting the orientation between two units, two populations of the CTD dimer interface are observed.

In contrast to the pliability of the RSV CA hexamer, the CA pentamer represents a rigid building block that acts as the local organization center that defines hexamer shape and, depending on the position on the lattice, overall core architecture.

It is therefore tempting to speculate that the increased deformability of hexamers in RSV, different to what has been previously found in HIV-1, could provide an explanation for the increased structural variability of cores in RSV. For example, when compared to RSV, the ∼9Å resolution structure of HIV-1 pentamer from authentic viral cores [9] does not reveal any deviation of its surrounding hexamers from true 6-fold symmetry (Fig. S8a) suggesting an overall lower flexibility of the HIV-1 hexameric CA unit in the mature assembly. Cone-shaped cores, as predominantly found in HIV-1, are determined by their distribution of pentamers, with 7 pentamers on the wide and 5 pentamers on the narrow end. An increased rigidity of HIV-1 hexamers could result in accumulated strain and tension around pentamers, resulting in their non-random distribution on mature cores, where pentamers locate to positions of defined curvature at the cone ends. This geometrical limitation could result in the more homogenous core size and shape distribution in HIV-1. In contrast, the increased flexibility of the RSV hexamer allows a more random distribution of pentamers in mature RSV cores. This enables RSV CA to even form inter-pentameric contacts, which can be regularly observed on polyhedrons and in particular T=1 icosahedral particles. No such pentamer-pentamer contacts have been yet observed for HIV-1 indicating that the described geometrical limitations in HIV-1 might render such interfaces unfavorable.

Further work on authentic CA cores or pleomorphic mature VLPs for other retroviruses will be required to determine the full spectrum of intra- and inter-hexameric CA interactions and if the rigidity of the pentamer as organizing center is a conserved feature of mature retroviral CA assemblies. Taken together, these findings shed light on the role for IP6 in the regulation of mature RSV assembly and also on the structural principles underlying retroviral mature core formation and shaping.

## Methods

### Protein purification and *in vitro* assembly

GagΔMBDΔPR and CASPNC proteins were purified using the SUMO-tag system [27] as previously described in [14]. The resulting SUMO purified proteins have an ectopic serine residue, which we previously reported does not influence assembly [13,15,28]. RSV CA protein was expressed in *E. coli* and purified using standard affinity and size exclusion chromatography described briefly hear. Bacterial pellets were resuspended in buffer (50 mM Tris-HCl pH 8, 2 mM TCEP), lysed by sonication, and the lysate cleared by ultracentrifugation. Nucleic acid was precipitated by the addition of 0.03% (v/v) polyethyleneimine followed by centrifugation. Ammonium sulfate was added to ∼20% saturation of the supernatant, and the precipitate was pelleted by centrifugation. The pellet was resuspended in buffer (50 mM Tris-HCl, 2 mM TCEP), and flowed through tandem QFF and SP columns (GE). CA protein was present in the flow through, and was ∼95% pure. Flow through was subjected to SEC through a Superdex 75 column (GE). Peak fractions corresponding to the CA protein were concentrated to 15-20 mg/mL, flash frozen in liquid nitrogen, and stored at −80.

*In vitro* assembly of GagΔMBDΔPR was performed by dilution assembly as previously described [29]. CASPNC was assembled via dialysis described briefly here. 30 µL containing 50 µM protein in storage buffer (20 mM Tris-HCl pH 8, 500 mM NaCl, and 2 mM TCEP) supplemented with ∼5 µM GT25 oligo, IP6 (if present) was dialyzed against assembly buffer (20 mM MES pH 6.2, 100 mM NaCl, 2 mM TCEP, IP6 (if present)) at 4° C for 4 hrs. GagΔMBDΔPR and CASPNC were spotted onto EM grids, negative stained with uranyl acetate, and imaged via TEM. *In vitro* assembly of CA with IP6 and NaPO_4_ was done by dilution assembly. Protein was diluted into assembly buffer (20 mM Tris-HCl pH 8, 2 mM TCEP) supplemented with NaPO_4_ or IP6, to final concentration of 290 µM. Samples were incubated at 22° C for 30 min and stored at 4° C until spotted onto TEM grids.

### Cell culture, virus release, and infectivity

HEK293FTs and IPPK-KO cells were maintained and plated for transfections and transductions as previously described [13]. RSV particles for *in vivo* assays were produced and transduced in a similar manner as HIV as previously described [13]. Flow cytometry gating strategy is shown in Fig. S9. Western blots and their analysis performed as previously described, and briefly here [13]. A rabbit anti-RSV-capsid antibody (NCI-Frederick, NCI 8/96) was used at a 1:500 dilution in 5% nonfat dry milk in PBS-tween for the 1 hr primary application at room temperature. A goat anti-Rabbit peroxidase conjugated antibody (Sigma, A0545) was used at a 1:10,000 dilution in 5% nonfat dry milk in PBS-tween for the 1 hr secondary application at room temperature. Blot images were converted to 8-bit, Fiji’s (ImageJ) gel analysis tools were used to calculate blot densities, and values exported to Excel for subsequent analysis in RStudio.

### Single particle cryo-EM

For cryo-EM 500 µM CA was assembled in 1 M NaPO_4_ at pH 8. The assembly reaction was diluted 1:4 to 250 µM NaPO_4_, and 3 µL were spotted onto glow discharged (45 sec, 20 mA) 2/2-3C C-flat grids. Samples were vitrified in liquid ethane using a Vitrobot Mark 4 (blot time of 2.5 sec and a blot force of 0). Imaging was done at 200 kV on a Thermo Fisher Talos Arctica TEM equipped with a BioQuantum post column energy filter and Gatan K3 direct detector using the SerialEM software package [30]. The data was collected with a nominal magnification of 63,000x in super resolution mode with a physical pixel size of 1.25Å/pixel. 50 frames were captured as movies with a total dose was ∼50e-/Å2. Image processing was done in RELION 3.1 [31], and software was maintained by SBGrid [32]. Specifically, motion correction was done using MOTIONCOR2 [33] in the RELION wrapper. CTF estimation and correction was performed using GCTF [34] in the RELION wrapper. For T=1 structure determination, 2394 micrographs were taken, from which 374 particles were manually picked and 2D classified to generate templates for auto-picking. Two rounds of auto-picking and 2D classification resulted in 150599 extracted particles. These particles were refined against EMD-5772 [24] (low pass filtered to 60Å) to a resolution of 4.6Å. Further iterative refinement, 2D and 3D classification, CTF refinement and Bayesian particle polishing resulted in a final set of 21498 particles that refined to a resolution of 3.1Å. T=3 VLPs were selected from the same initial 2D classification job to generate templates for auto-picking. Three rounds of auto-picking and 2D classification resulted in 2309 particles. Subsequent rounds of 2D and 3D classification resulted in a set of 599 particles that refined against EMD-5773 [24] (low pass filtered to 60Å) to 8.6Å. 2D and 3D classification, CTF refinement, and Bayesian particle polishing, resulted in a set of 406 particles that refined to 7.6Å.

### Cryo-electron tomography

Cryo-electron tomography sample preparation and data collection were done essentially as described previously [7,14]. 10nm colloidal gold was added to CASPNC VLPs and 2.5 µl of this solution was then applied to degassed and glow discharged 2/2-2C C-flat grids. The samples were vitrified in liquid ethane using a Vitrobot Mark 4 (blot time of 2s, wait time of 5s, blot force 0) and stored in liquid nitrogen until imaging.

Tilt series were acquired on an FEI Titan Krios, operated at 300 keV, equipped with a Gatan Quantum 967 LS energy filter and a Gatan K2xp direct electron detector using the SerialEM software package [30]. The slit width of the filter was set to 20 eV. Areas of interest for high-resolution data collection were identified in low magnification montages. Prior to tomogram acquisition, gain references were acquired and the filter was fully tuned. Microscope tuning was performed using the FEI AutoCTF software [35].

The nominal magnification was 105,000x, resulting in a pixel size of 1.328 Å/pixel. Tilt series were acquired using a dose-symmetric tilt-scheme [36], with a tilt range from 0° to −66° and +63° in 3° steps. Tilt images were acquired as 8k x 8k super-resolution movies, consisting of 10 frames. The dose rate was set at ∼ 2.5 e^-^/Å/sec and aiming for a target dose of ∼156 e^-^/Å^2^/tilt series. Tilt series were collected at nominal defocus between −1.5 and −4 µm. Data acquisition information is also provided in Table S1.

### Cryo-ET image processing

K2 super-resolution movies were aligned on-the-fly during data acquisition using the SerialEMCCD frame alignment plugin and tilt series were automatically saved as 2x binned mrc stacks. Prior to further processing, tilt series were sorted and bad tilts (e.g. images with large shifts or objects blocking the beam) were removed using MATLAB scripts.

CTF-estimation was performed using CTFFIND4 (version 4.1.10) [37] on each tilt individually. Images were low-pass filtered according to their cumulative electron dose, as described previously [38]. Tilt series alignment of the exposure-filtered micrographs was performed in the IMOD software package [39] and 3D-CTF-corrected tomograms were reconstructed by weighted back-projection using novaCTF [40]. The CTF correction method was multiplication with astigmatism correction and a Z-slab thickness of 15 nm. The resulting tomograms were consecutively binned 2, 4, and 8 times via Fourier cropping.

In total 49 tomograms were used for further processing. Subtomogram averaging was performed in Dynamo [41]. Additional intermediate processing steps and steps for context-based classification (as described below) were done using MATLAB scripts, employing functions from Dynamo, AV3 [42] and TOM [43] software packages. Complete statistics for the sizes of the different datasets are given in Table S2.

#### Subtomogram averaging of regular CASPNC tubes

In order to determine the structure of the CA hexamer from regular tube VLPs, subtomograms extracted along the tubular surface were subjected to iterative averaging and alignment, as previously described [14]. Briefly, initial subtomogram positions and Euler angles were set based on tube radius and splines fitted to the respective tubular axes using MATLAB and 3dmod software [39]. Initially, a single tube, derived from a tomogram with a nominal defocus of −2.5 µm was chosen to derive a *de novo* starting reference at bin8. This reference was then used for the alignment of the entire tube VLP dataset consisting of 236 tubes.

Bin 8 starting extraction positions on tube surfaces were oversampled ∼4 times. After the first round of bin 8 alignment, all subvolumes that had converged onto the same position of the tubular lattice were removed using a subvolume-to-subvolume distance cut-off. Subvolumes that contained no protein density or did not align against the reference were removed based on a cross-correlation (CC) threshold.

In total, 10 rounds of alignment were performed, progressively reducing the angular search range, adapting low-pass filters and reducing binning. At every unbinning step (i.e. going from bin8 to bin4) subtomograms were re-extracted from the respective unbinned tomograms at positions determined in the previous binned alignments. New averages were then generated using Euler angles determined in the previous binned alignment round. C2 symmetry was applied throughout the entire processing of this tubular dataset.

At bin2, data was split into even/odd half-sets, which from hereon were treated completely independently. Up to bin2 alignment low-pass filter settings were restricted to 32 Å, ensuring that even/odd datasets were not aligned on identical features beyond this resolution.

Final bin1 averages displayed varying anisotropy due to a preferential orientation of the tubes in the tomograms with respect to the tilt axis. To compensate for the anisotropy in the averages, weighted averaging of the 1x binned data was performed using modified wedge masks [14,44]. Mask-corrected Fourier-shell correlation (FSC), using gaussian-filtered masks, was employed to determine the resolution at the 0.143 criterion. The half maps were then combined, sharpened and filtered to the measured resolution [45].

#### Lattice annotation of polyhedrons

In order to initially annotate the CA lattices from all VLPs (in particular irregular polyhedrons, but also again tubes), an in-house written MATLAB script for template matching was used, in part employing functions from the Dynamo [41], AV3 [42] and TOM [43] software packages. Based on previous observations of mature retroviral CA assemblies [3,9], the CA lattice of RSV CASPNC VLPs was assumed to consist of C2-symmetric CA hexamers (as seen in our regular tube VLPs), C6-symmetric CA hexamers and C5-symmetric CA pentamers (in polyhedral VLPs). Therefore, 3 different templates following these symmetries were generated in a similar fashion as described above for the C2-symmetric hexamer, but with the following differences. Subvolumes were first aligned in bin 8. Subsequently, three additional bin4 alignment rounds allowed sorting of subtomograms into pentamers (C5) and hexamers (C2 and C6), to generate three new averages for these CA assembly types, which were then used as templates for template matching on bin8 tomograms.

For template matching, a cylindrical mask encompassing the central CA pentamer or hexamer and one ring of adjacent hexamers was applied to the template. Both the reference and tomogram were bandpass-filtered (425Å-30Å) before CC calculation. Phi and non-Phi angles were scanned over a range of 360° in 10° and 15° steps, respectively. To select the highest scoring cross-correlation peaks from the resulting cross-correlation array, a lattice connectivity analysis was performed. The following criteria were used to asses pairs of local cross-correlation maxima determined for hexamers/pentamers to define a maximum-connectivity network (which is forming a distinct VLP): minimum spacing between hexamers/pentamers = 60 Å, maximum spacing between hexamers/pentamers = 110 Å, minimum curvature (i.e. difference between the tangential plane and a vector connecting the respective pair of peaks) = −15°, maximum curvature = 40°, maximum difference of normal vectors corresponding to the respective CC peaks = 45°. Only networks containing >20 cross-correlation peaks for either of the three used templates were considered as VLPs and used for subsequent steps.

#### Context-based classification of CA subtomograms

The irregularity of the CA lattice in polyhedron CASPNC assemblies limited the achievable resolution using conventional subtomogram averaging (as described for the regular tube VLPs). To overcome this limitation, subvolumes were classified according to their local geometrical context. Specifically, pairs of adjacent pentamers/hexamers (so called unit pairs, where a unit refers to a pentamer or hexamer, resulting in possible unit pairs of hexamer/hexamer, hexamer/pentamer or pentamer/pentamer) were sorted into classes based on their local lattice arrangement.

To this end two classification criteria were used, based on the following observations of our data:

i. pentamers within polyhedrons and hexamers from tubes are regular objects with defined C5 and C2 symmetry, respectively (Fig 2a and b);
ii. the hexamers in the vicinity of a pentamer have varying geometry (Fig. S5);
iii. hexamers, which are further distant from a pentamer, adopt a more tube-like geometry with C2 symmetry.

The first criterion for dimer classification is the position of pentamer(s) in a given unit pair. Each hexamer adjacent to a pentamer has all of its dimeric connections numbered relative to the position of the pentamer (Fig. S3a), resulting in numbers 1’ to 6’ for the first unit (i.e. unit A), and 1’’-6’’ for the second (i.e. unit B). Number 1’ or 1’’ indicates that one unit from the dimer is pentamer. Numbers 2’ (which is the same as 6’’) or 6’ (which is the same as 2’’) indicate that a pentamer neighbor is shared between unit A and B. Numbers 3 to 5 indicate a pentamer that is directly in the vicinity of only one unit of the pair.

The second criterion for dimer classification is the orientation of the unit pair with respect to the tube geometry (Fig. S3b). In a regular tube each hexamer is surrounded by six C2 hexamers (except for the tube ends). There are therefore 3 distinct orientations of two neighboring hexamers with respect to the tube geometry. This criterion was used for classification of hexamer-hexamer dimers, in which at least one did not neighbor with any pentamer.

Using these criteria, unit pairs were sorted into 6 groups, which were then further divided into sub-classes (Fig. S3c). Group 1 comprises dimers, in which one or both of the respective units are pentamers (in positions 1’ and 1’’). For the initial consensus alignment, any further pentamers in positions 2 to 6, were not considered during the classification. For the final round of consensus alignment, however, additional surrounding pentamers in positions 2-6 (with respect to the hexamer) were used to sub-classify the hexamer-pentamer connections. Group 2 comprises two neighboring hexamers, which share one or two pentamer neighbors in positions 2 and 6. Similarly, as in the previous group, further pentamers in positions 3’ −5’ are only considered in sub-classes in the final round of consensus alignment. Group 3 encompasses dimers of hexamers, in which each hexamer has exactly 1 pentamer neighbor in positions 3, 4 or 5. Group 4 contains dimers, in which only one hexamer has a pentamer neighbor (positions 3, 4, or 5). Since the geometry of the second hexamer is not constrained by a pentamer in its vicinity, the dimer orientation with respect to the tubular geometry is used to further sort particles (Fig. S3c). Group 5 contains dimers of hexamers without pentamers in position 2 to 6 (i.e. in regular CASPNC tubes not having pentamers). Again, classification according to the orientation with respect to the tube geometry was used to yield 3 sub-classes. Group 6 comprises dimers, in which hexamer A contains pentamers in two out of three positions: 3’,4’ and 5’, and hexamer B does not have any pentamer in its vicinity. The angle between the dimer and the tube axis is in this case neglected, since further classification would yield very sparsely populated classes. This context-based classification laid foundation for the consensus alignment, as specified below.

#### Consensus alignment of tubular and polyhedral CASPNC VLPs

Instead of refining the positions and orientations of isolated hexamer and pentamer units, an alignment of unit pairs classified into one of the 6 groups was performed. To this end, for each unit pair a set of alignments, always masking the central and one of the adjacent units, was performed. Hence, the number of alignments *N* per unit corresponded to the number of its directly surrounding units in the lattice, yielding *N*(x,y,z) coordinates and *N*(*φ,ϑ,ψ*)Euler angle triplets. The *N* set of alignments was then used to determine a consensus alignment for a unit depending on its neighbors. The consensus position of the unit center was calculated as weighted mean of the (x,y,z) coordinate sets.

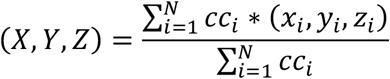

Where *x*_*i*_,*y*_*i*_,*z*_*i*_ and *X,Y,Z* are the positions of a given particle in Cartesian coordinates for different measurements and their consensus, respectively, and *cc*_*i*_ is cross-correlation.

The consensus orientation was calculated as the weighted sum of the normal vectors corresponding to the respective Euler angle triplets (*φ,ϑ,ψ*).

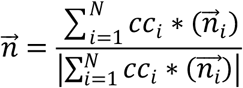

Where 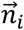 and 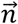 are normal vectors obtained for different measurements and their consensus, respectively. The cross-correlation against the respective classes was used as a weighting function.

An example of the consensus alignment for a hexamer in the lattice of a polyhedron is given in (Fig. S4b).

For the alignment the coordinates determined using template matching in bin8 tomograms were first unbinned to bin4, and subjected to 2 rounds of (standard) subtomogram alignment. 4 and 2 rounds of consensus alignment classified to groups 1 to 6 were performed with bin4 and bin2 data, respectively. One final round of consensus alignment with the group sub-classes was then performed with bin2 data.

The context-based classification and consensus alignment also provided information on the CA lattice geometry in polyhedrons, specifically the tilt, twist and distance geometry of unit pairs in the different classes (Fig. S10). This information was utilized to further refine the unit positions in each class based on the mean geometry of the lattice in each class. To this end, the coordinates and Euler angles of each unit pair were optimized to minimize tilt, twist and distance deviation of each unit pair from the mean value of the respective class.

This geometry optimization was performed before and after the last round of consensus alignment on 2x binned data, and resulted in improved averages and slightly increased resolution, compared to the consensus alignment alone.

As in the case of processing of the C2-hexamer from tubes, the data were split into half-sets at the bin2 stage. Specifically, in this case, the data were split into two halfsets of comparable size on the VLP level, in order to ensure complete separation of the subvolumes. Again, low-pass filters were set no lower than 32 Å prior to this step, to avoid aligning on higher resolution features within the unsplit dataset.

In order to generate the final class averages, 2 rounds of conventional subtomogram alignment were performed on unbinned data using the translational and orientational parameters from the previous binned consensus alignments. At this step, boxes were centered at the dimer interface and hence no more consensus refinement was performed.

### Data visualization, model building and analysis

Cryo-EM and cryo-ET data and structures were visualized in IMOD [39] and UCSF Chimera [46]. Visualization of VLP lattice maps was done using the PlaceObject plugin in UCSF Chimera [12].

The resolution of our electron microscopy maps for the C2-symmetric hexamer obtained from the regular CASPNC tubes and the CA pentamer from T1 particles allowed us to refine existing models of RSV CA (pdb 3TIR) [22]. At the obtained resolution for the C2-symmetric hexamer (4.3 Å), the helical pitch was visible and also several large side chains (e.g. tryptophans, phenylalanines, arginines or lysines) could be identified. Small side chains and also negatively charged side chains were not clearly visible at this resolution. Therefore, rotamer refinements of residues were not possible.

The pseudo atomic model of RSV CA (pdb 3TIR) was used as a starting model for refinement into the T1 particle map. The CA_NTD_ and CA_CTD_ of one CA monomer of pdb 3TIR were independently placed into the EM-density using the rigid body fitting option in UCSF Chimera. Subsequently the linker connecting the two CA domains was joined in Coot. To account for the different monomer-monomer interactions in the RSV CA pentamer, the monomers were replicated according to the inherent 5-fold symmetry of the map and an additional ring of CA-_CTD_’s was rigid-body fitted into the EM-densities of the surrounding CA pentamers. A map segment (defined by a mask extending 3 Angstrom around the rigid body fitted model) was extracted, and real-space coordinate refinement against the EM-density was performed using Phenix [47]. This was iterated with manual model building in Coot [48], similar as described previously [14]. In brief, secondary structure restraints and non-crystallographic symmetry (NCS) restraints were applied throughout all refinements. Each Phenix iteration consisted of 5 macro cycles, in which simulated annealing was performed in every macro cycle. Atomic displacement parameter (ADP) refinement was per formed at the end of each iteration.

For modeling the mature CA assembly in the C2-symmetric hexamer in regular CASPNC tubes, one CA monomer of pdb 3TIR was used. The monomer was rigid body fitted into the EM density three times to accommodate the 3 symmetry independent copies of CA in the C2-symmetric hexamer. The residues in the first strand of the N-terminal β–hairpin were removed, as no ordered density corresponding to these residues was present. As symmetry-independent monomers show differences in their respective orientations of their CANTD and CA_CTD_, the fit was further manually refined in Coot. Subsequently, the symmetry independent monomers were expanded according to the C2 symmetry of the cryo-EM density, completing the hexameric CA assembly. An additional ring of CA_CTD_’s was rigid-body fitted into the EM-densities of the surrounding CA hexamers, to shield the inner hexameric ring. Refinement was then performed similarly as described above, iterating between automatic refinement in Phenix and manual model building in Coot.

The model of the CA monomer derived from real-space refinement into the T1 particles was used to generate pseudo-atomic models of the hexameric and pentameric CA assembly in the CASPNC polyhedrons. CA_NTD_ and CA_CTD_ domains were separately fitted into the respective maps using the rigid-body fit functionality in UCSF Chimera. For CASPNC polyhedron-reconstructions no further real-space refinement of the models was performed.

The quality of the pentamer CA model derived from T1 particles and the C2 symmetric hexamer CA model from regular tube VLPs was validated using MOLPROBITY [49].

All comparisons and RMSD calculations were performed in UCSF Chimera or Matlab, between the C-alpha backbone atoms of selected residues.

## Acknowledgements

This work was funded by the National Institute of Allergy and Infectious Diseases under awards R01AI147890 to R.A.D., R01AI150454 to V.M.V, R35GM136258 in support of J-P.R.F, and the Austrian Science Fund (FWF) grant P31445 to F.K.M.S. Access to high-resolution cryo-ET data acquisition at EMBL Heidelberg was supported by iNEXT (grant No. 653706), funded by the Horizon 2020 program of the European Union (PID 4246). We thank Wim Hagen and Felix Weis at EMBL Heidelberg for support in cryo-ET data acquisition. This work made use of the Cornell Center for Materials Research Shared Facilities, which are supported through the NSF MRSEC program (DMR-179875). This research was also supported by the Scientific Service Units (SSUs) of IST Austria through resources provided by Scientific Computing (SciComp), the Life Science Facility (LSF), and the Electron Microscopy Facility (EMF).

## Tables

**Table S1:**
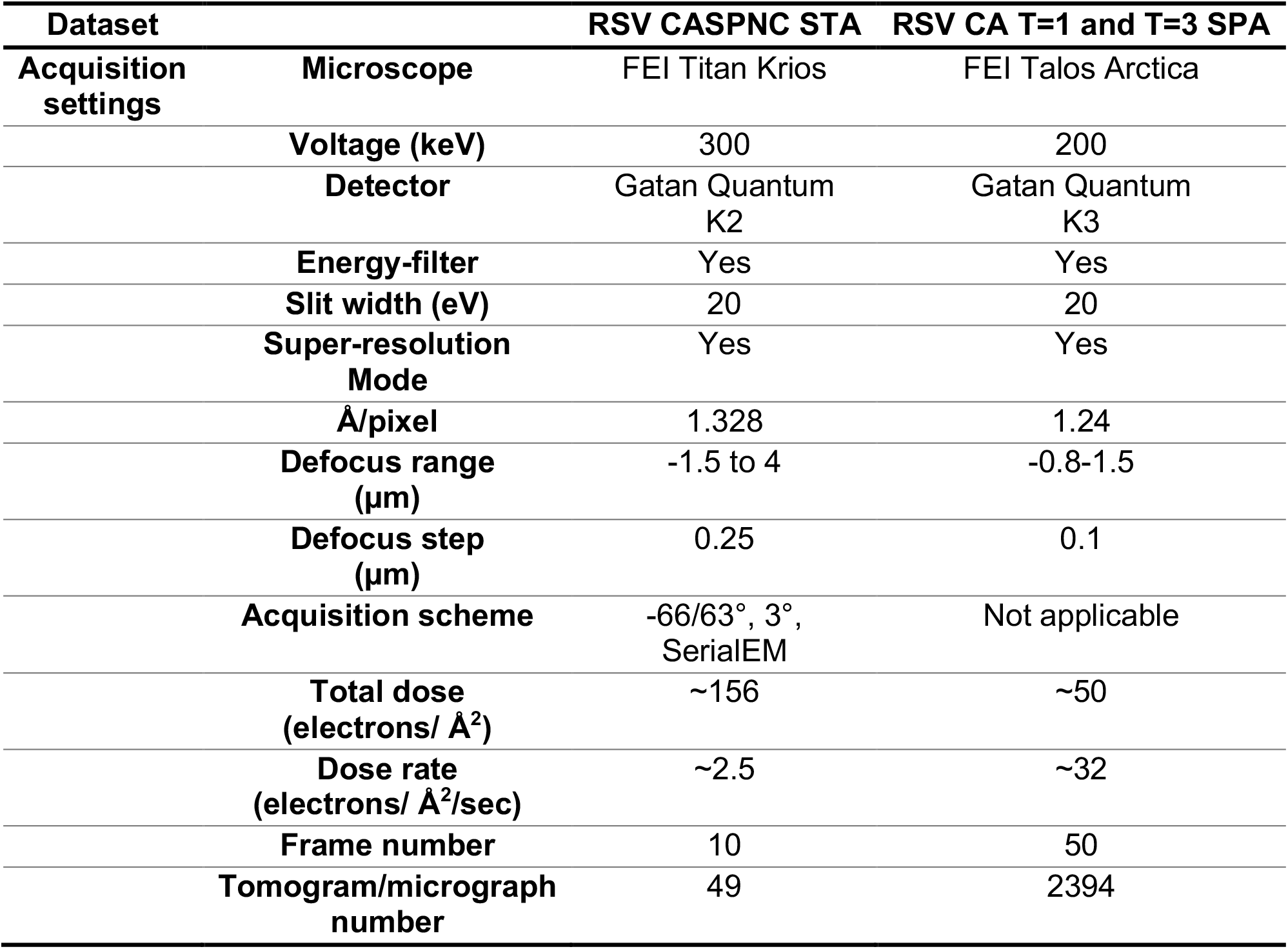
Data acquisition statistics.

**Table S2:**
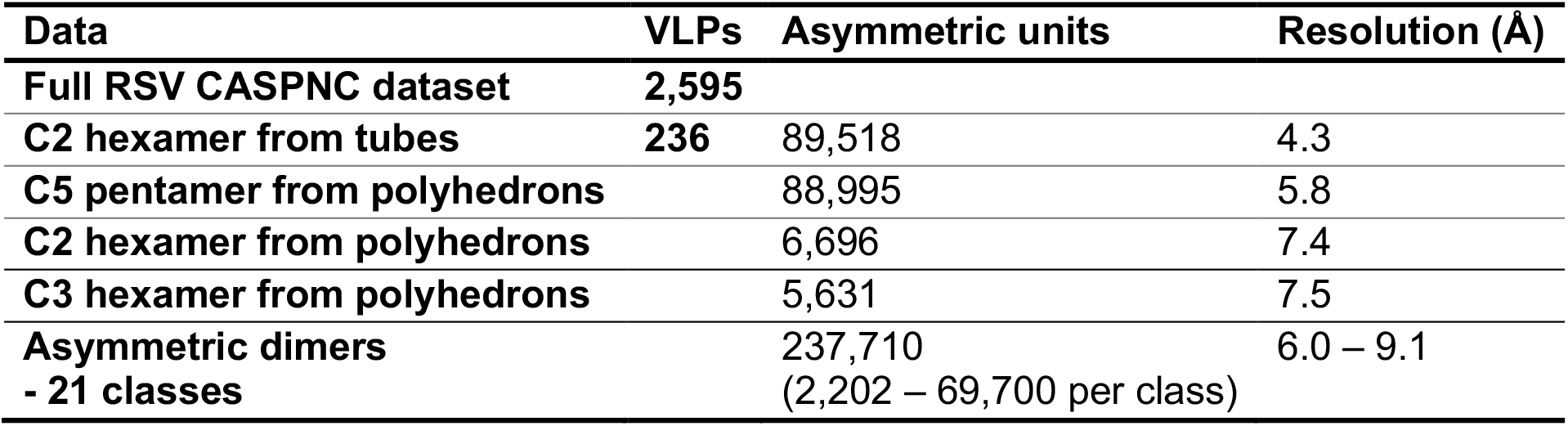
Data set statistics.

## Supplemental figures

**Figure S1:**
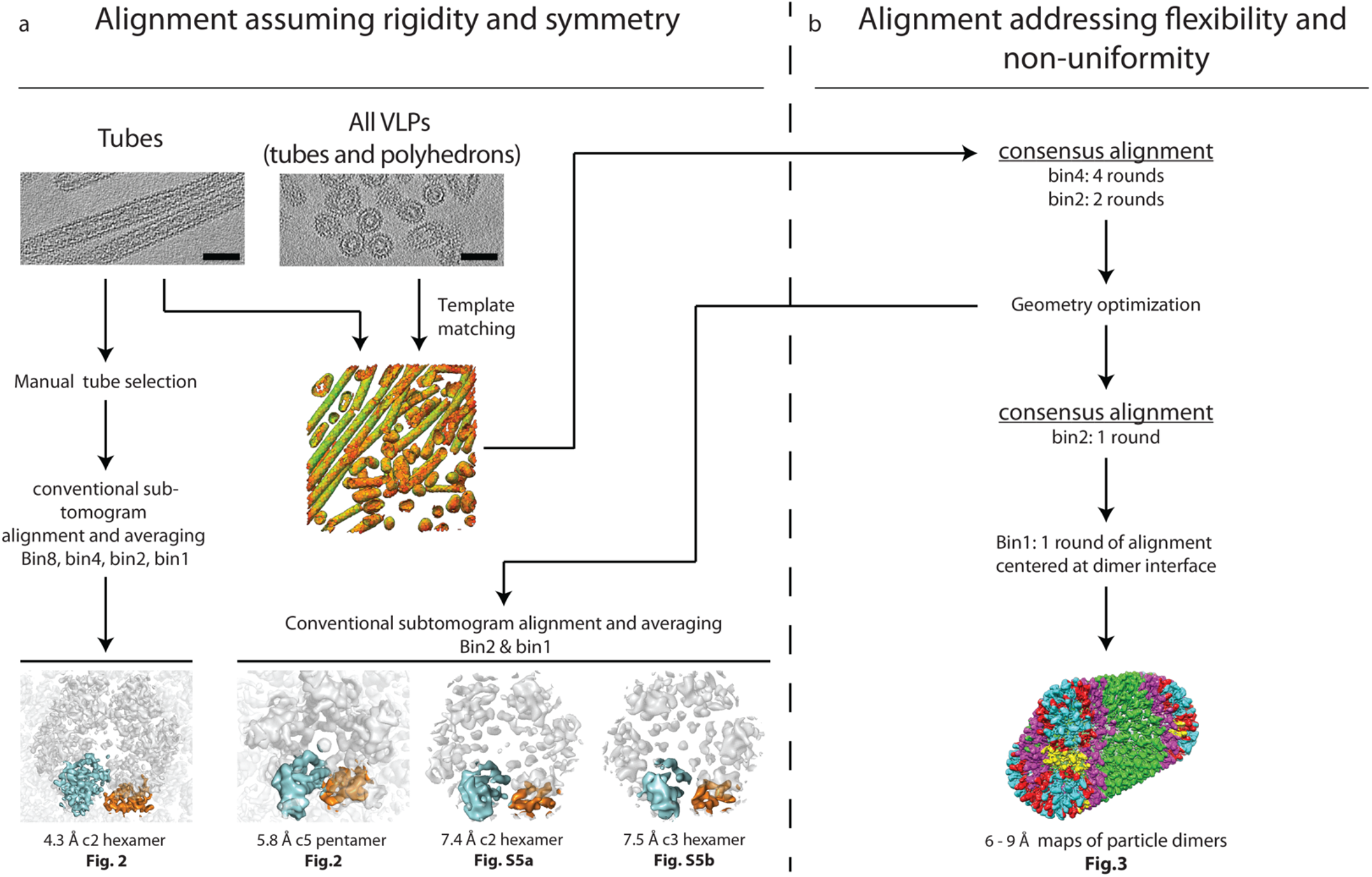
Subtomogram averaging workflows. Basic schematic summary of the subtomogram averaging workflows employed in this study. **a)** On the left side the standard subtomogram alignment assuming rigidity and symmetry is shown. **b)** The parts of the workflow using the novel consensus alignment routines are shown on the right side. Connections between the two approaches, underlining their modularity, are annotated by arrows. Details on the novel alignment and classification strategy are given in *Materials and Methods* and Figure S3.

**Figure S2:**
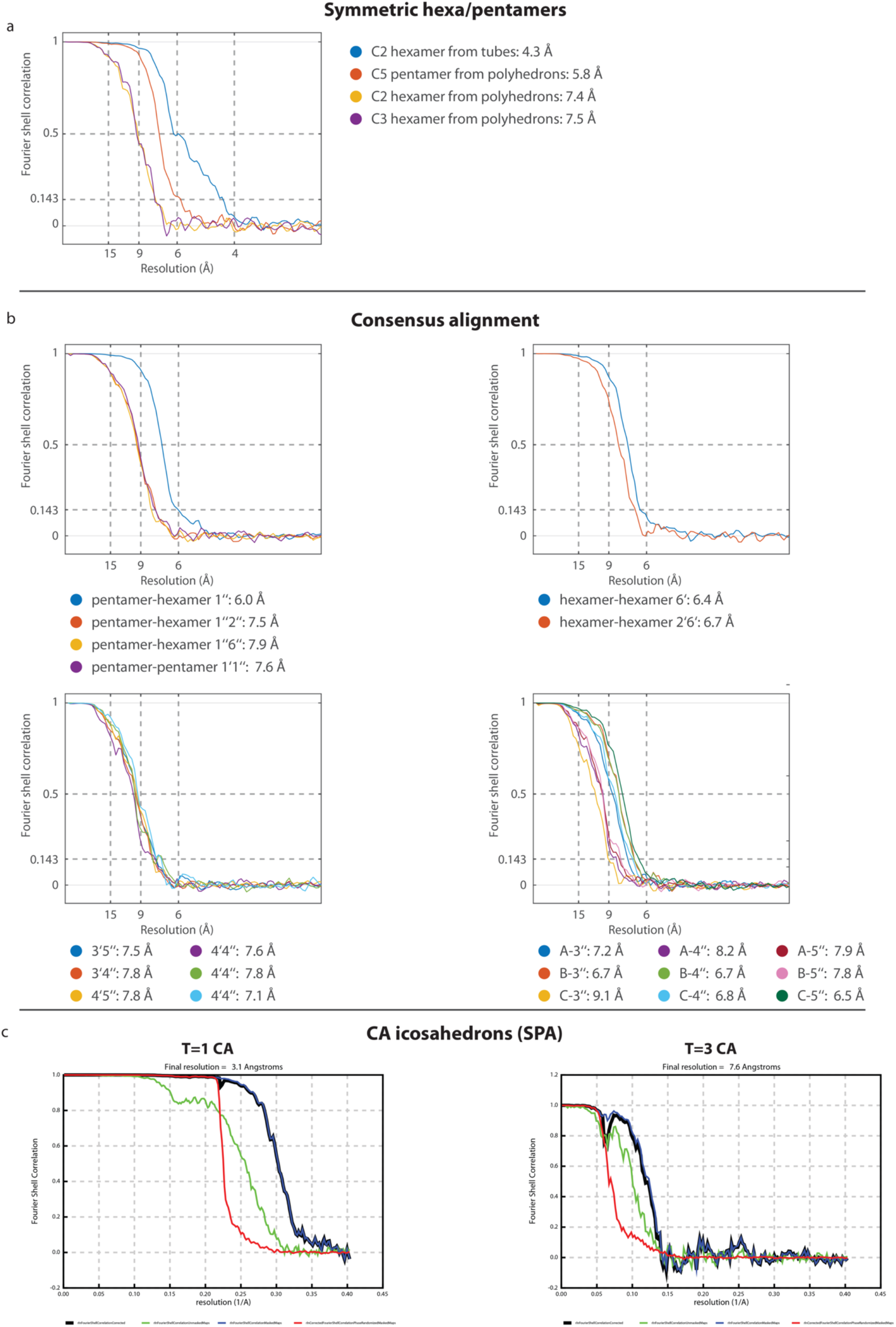
Resolution estimation via Fourier Shell correlation. a) FSC curves for pentamers and hexamers solved by subtomogram averaging, while utilizing the corresponding symmetries (C2, C3, C5). **b)** FSC curves for different classes from groups I-IV, which were analyzed using consensus alignment. **c)** FSC curves for RSV CA icosahedrons analyzed by single-particle analysis.

**Figure S3:**
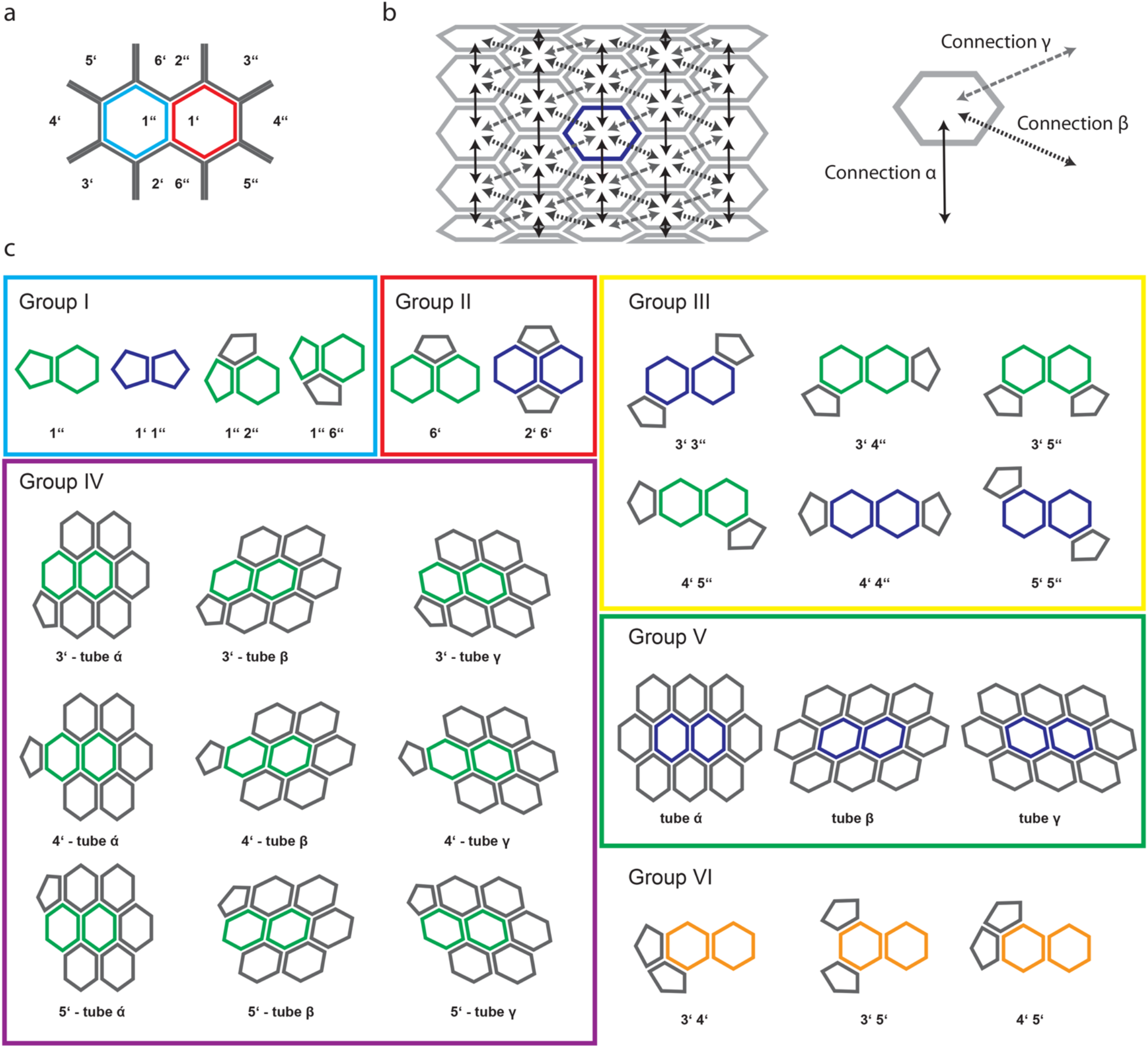
Context-based classification. Graphical summary of the basic principles underlying the context-based classification. **a)** The numbering scheme for classification. Numbering of a unit pair is derived from the position of pentamer with respect to the two units of a unit pair; the central unit is colored cyan and any pentamers neighboring with this unit are marked with a single prime; the adjacent unit of the pair is colored red and any neighboring pentamers are marked with a double prime. **b)** The three different hexamer-hexamer connections in a regular CASPNC tube are shown. The tube axis in the scheme is horizontal. Connections α, β, and γ are shown in solid black, short-dashed dark grey, and long-dashed light grey arrows, respectively. These hexamer-hexamer connections are used to subclassify Groups 4 and 5. **c)** Schematic depiction of different classes distinguished during context-based classification. Groups of classes with related context are highlighted by a colored frame, as in Fig. 3: cyan: group I; red: group II; yellow: group III; purple: group IV; green: group IV. Hexagons and pentagons represent CA hexamers and pentamers, respectively. Two hexamers and pentamers form a unit pair, a basic unit, which is a subject to classification. The unit pairs colored blue consist of two units with identical geometric context based on the classification criteria. The unit pairs colored green consist of two units with different geometric context based on the classification criteria, resulting in two classes that are related by a 180 degree rotation during consensus alignment. In case of the orange unit pairs from Group VI the context of one unit is neglected, as it would lead to too sparsely populated classes. Therefore, group VI was used only during consensus alignment, but not for generating the final bin1 maps.

**Figure S4:**
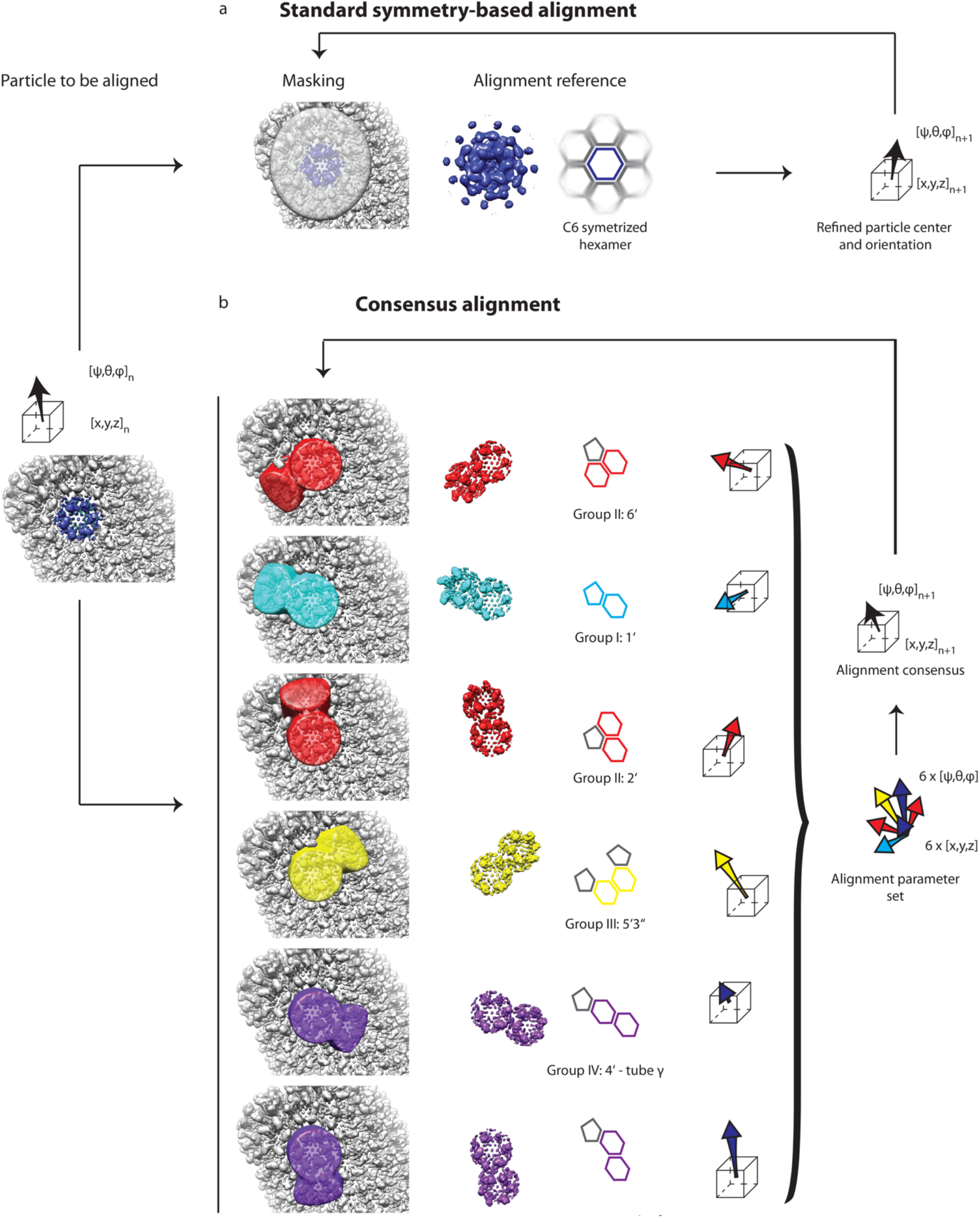
Graphical summary of the consensus alignment. a) Alignment of CA hexamers using the “standard” approach: from left to right – a single hexamer in the context of a grey VLP is highlighted in blue; the subvolume containing this hexamer is masked and iteratively aligned against a symmetrized reference, refining translation and orientation parameters of the subvolume. **b)** The consensus alignment builds on the same principles as the “standard” alignment routine. However, the alignment is performed for each unit multiple times equal to the number of its adjacent units. Each pair of the central and adjacent unit are classified according to their context (see groups in Figure S3), and a reference representing the respective class is used for alignment in each case, while applying a tight mask around the central and the respective adjacent unit. Therefore, in each round multiple alignment parameters per unit are obtained. The consensus of the individual alignments per unit yields the alignment parameters for the next iteration.

**Figure S5:**
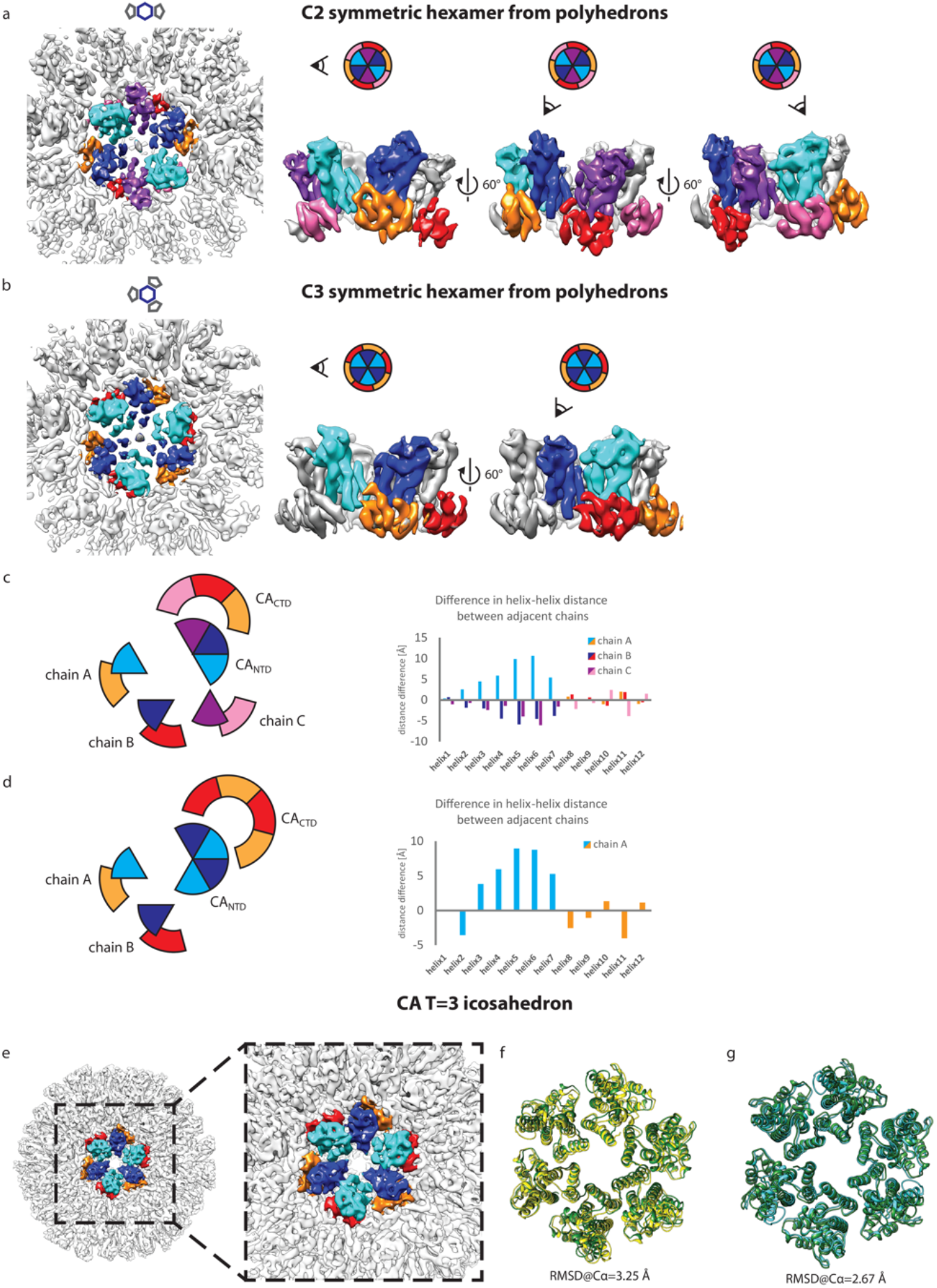
The RSV CA pentamer structurally distorts adjacent hexamers. **a**) Distortion of a CA hexamer flanked by two pentamers in polyhedral VLPs. Left: EM-density of the C2-symmetric hexamer surrounded by two pentamers, shown in top view; middle and right: side views showing neighboring symmetry-independent CA monomers. Symmetry-independent CA monomers are distinguished by colors: cyan/orange, blue/red, and purple/pink, respectively. The viewing angle is indicated by an eye symbol. The two hexamer CA_NTD_’s facing the pentamers show a larger opening. **b)** Distortion of a CA hexamer flanked by three pentamers in polyhedral VLPs. Views are identical to A). Side views are showing neighboring symmetry-independent CA monomers in cyan/orange, and blue/red, respectively. Again, the separation of the hexamer CA_NTD_’s facing the pentamers is clearly visible. **c-d)** Distortion analysis of C2-, and C3-symmetric CA hexamers, respectively. Left: schematic view of CA hexamer showing symmetry-independent CA copies. The color code is as in (a and b). Difference in distance between identical helices of two adjacent CA copies measured per helix and symmetry-independent CA copy. **e-g)** T=3 CA icosahedron. **e)** isosurface representation of a T=3 icosahedron solved by single particle analysis cryo-EM; the dashed line highlights one hexamer colored identically as in panel B, with three symmetry independent CA copies colored differently. **f)** The T=3 icosahedron hexamer CA model obtained via rigid body fitting of the individual CA domains separately (green) into the EM-density. The identical model is rotated by 60° (yellow) and shown in superimposition. The RMSD of 3.25 Å shows that the T=3 icosahedron hexamer is not 6-fold symmetric. **g)** Comparison of the CA hexamer model from T=3 icosahedrons (green) and of a C3-symmetric CA hexamer from polyhedrons made by rigid body fitting of the individual CA domains separately (cyan). The RMSD of 2.67 Å indicates that both models are distorted by the presence of adjacent pentamers, but the distortion is less apparent in the T=3 icosahedra.

**Fig. S6:**
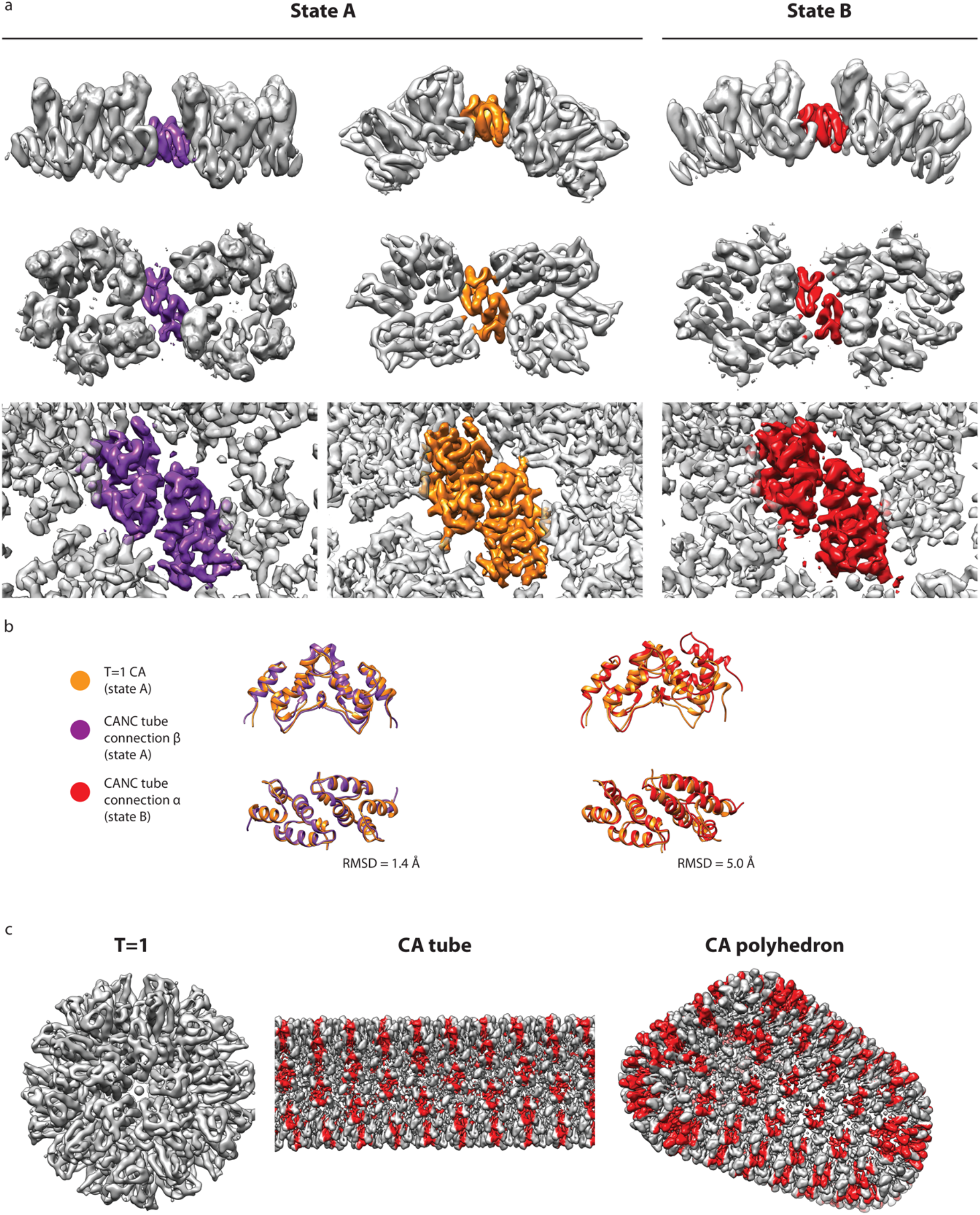
Two states of RSV CA_CTD_ dimer interface. **a**) Two distinct forms of CA_CTD_ dimer (as determined by clustering of pairwise RMSD measurements) were observed in RSV VLPs. State B is found only in high curvature hexamer-hexamer interfaces, whereas state A is found in pentamer-pentamer, pentamer-hexamer, and low-curvature hexamer-hexamer interfaces. Top and middle rows: cryoEM maps showing two adjacent hexa/pentamers filtered to 8 Å from side and top view, respectively; bottom row: cryoEM maps filtered to the resolution determined at 0.143 criterion. Two adjacent CA_CTD_s are colored according to the following color code: purple – tube connection β (hexamer-hexamer); orange – T=1 CA (pentamer-pentamer); red – tube connection α (hexamer-hexamer). **b)** Real-space refined models are overlaid in order to illustrate the difference between state A and state B dimer, and similarity of state A dimers found in different VLPs. Left: comparison of two state A CTD dimers from CASPNC tube connection β (purple), and T=1 CA (orange); right: comparison of state A CTD dimer from T=1 CA (orange) and CASPNC tube – connection α. **c**. The two states are highlighted on different shapes of RSV VLPs: grey – state A; red – state B.

**Figure S7:**
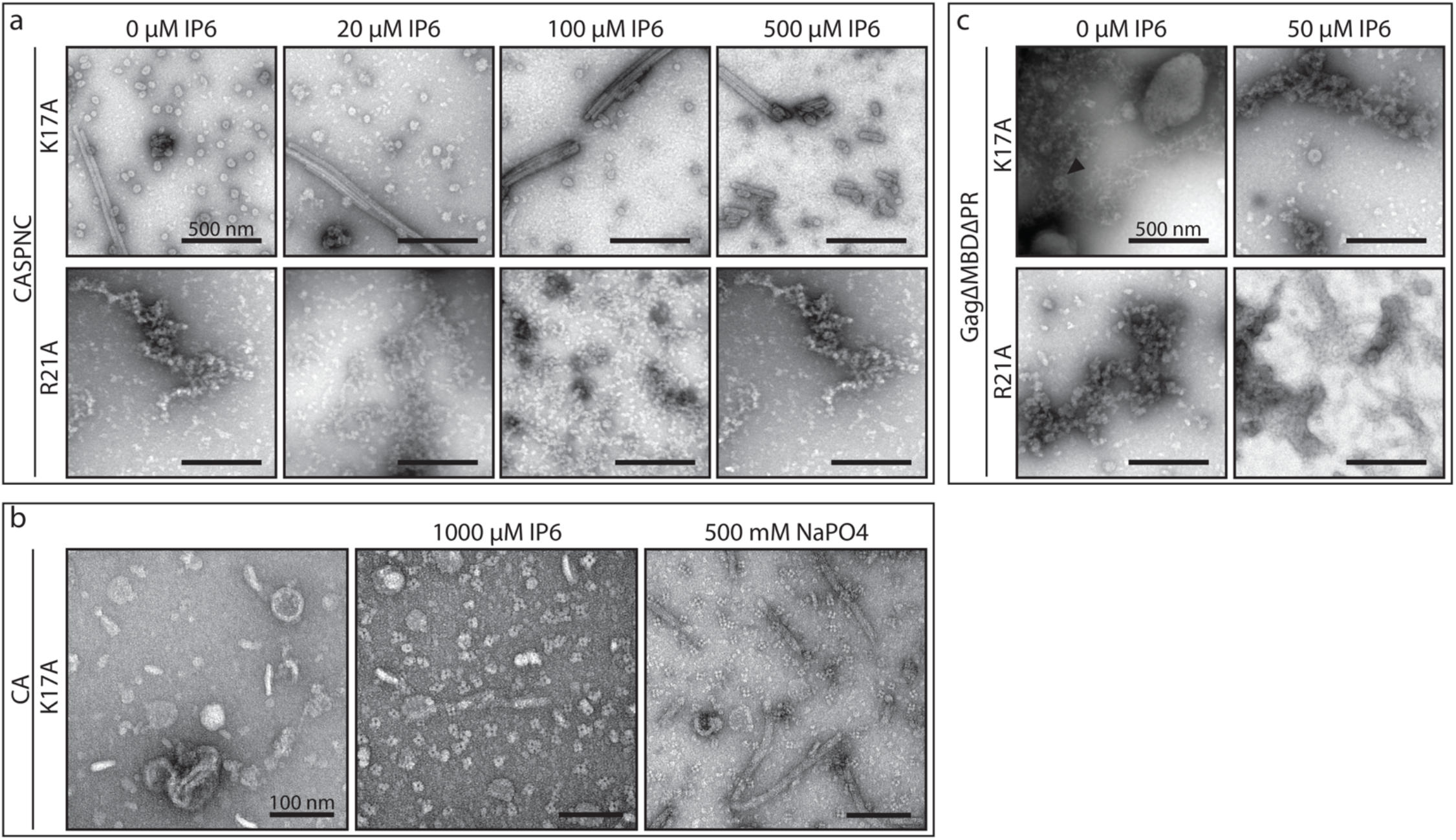
CASPNC K17A and R21A mutant assemblies. a) CASPNC K17A or R21A mutants assembled with increasing concentrations of IP6. **b)** CA K17A protein without and with IP6 or NaPO_4_. **c)** Assembly of GagΔMBΔPR K17A or R21A mutants without or with IP6

**Figure S8:**
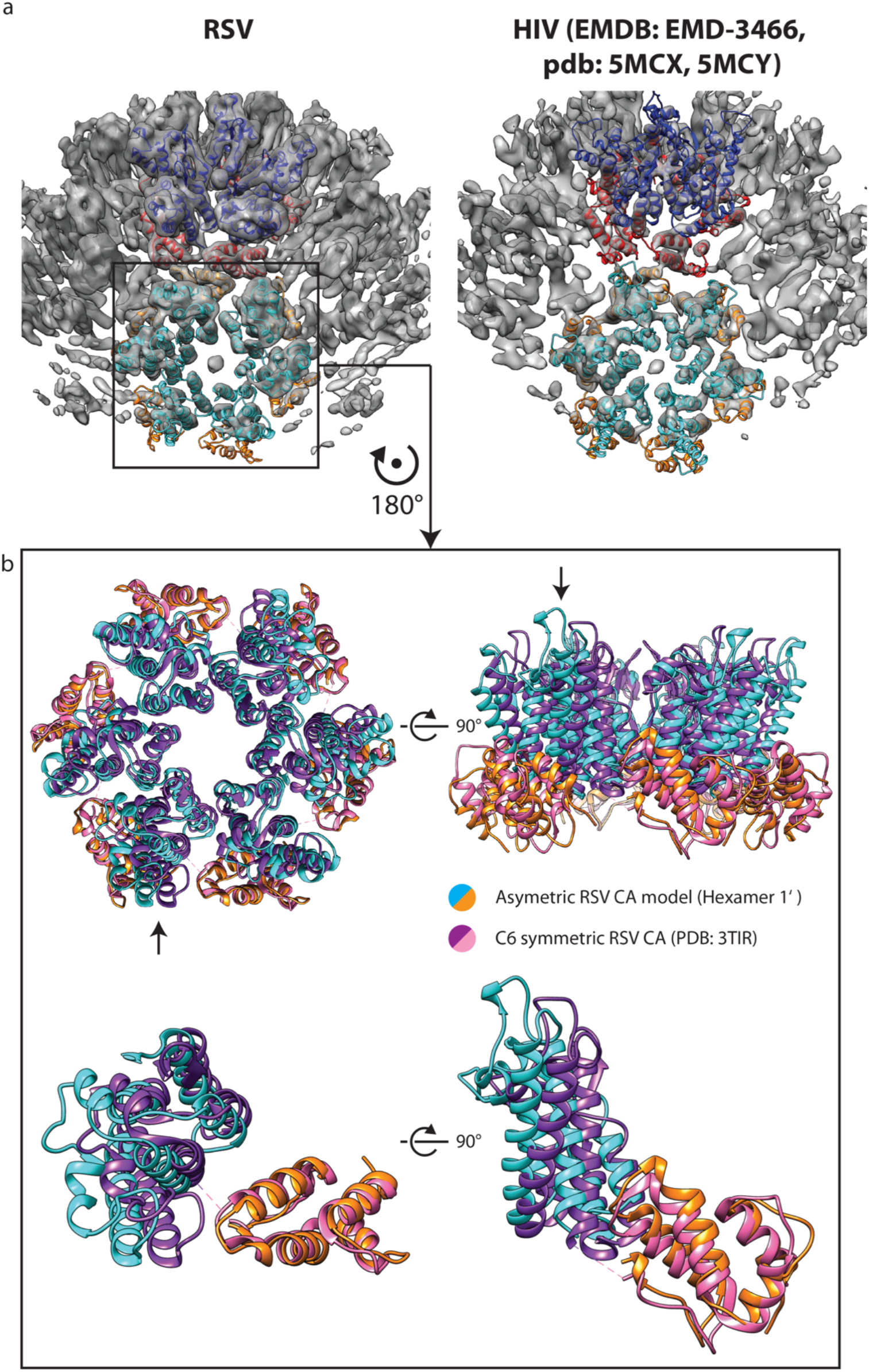
CA hexamer flexibility in RSV and HIV-1. a) Isosurface representation of the RSV (left) and HIV-1 (right) CA pentamer and its surroundings with models of the CA pentamer and CA hexamer fitted into the density (EMDB and PDB codes for the HIV-1 structure [9] and model are annotated). Note that the RSV CA pentamer induces an opening in the adjacent CA hexamer. **b)** Comparison of the models of RSV CA hexamer derived by subtomogram averaging and X-ray crystallography [22]. Cyan/orange – asymmetric model obtained by rigid body fitting of individual CA domains into the cryo-electron microscopy density of the hexamer adjacent to a pentamer; purple/pink – C6 symmetric X-ray crystallographic model (pdb 3TIR). The hexamer and one isolated monomer (annotated with an arrow) are shown in top and side views.

**Figure S9:**
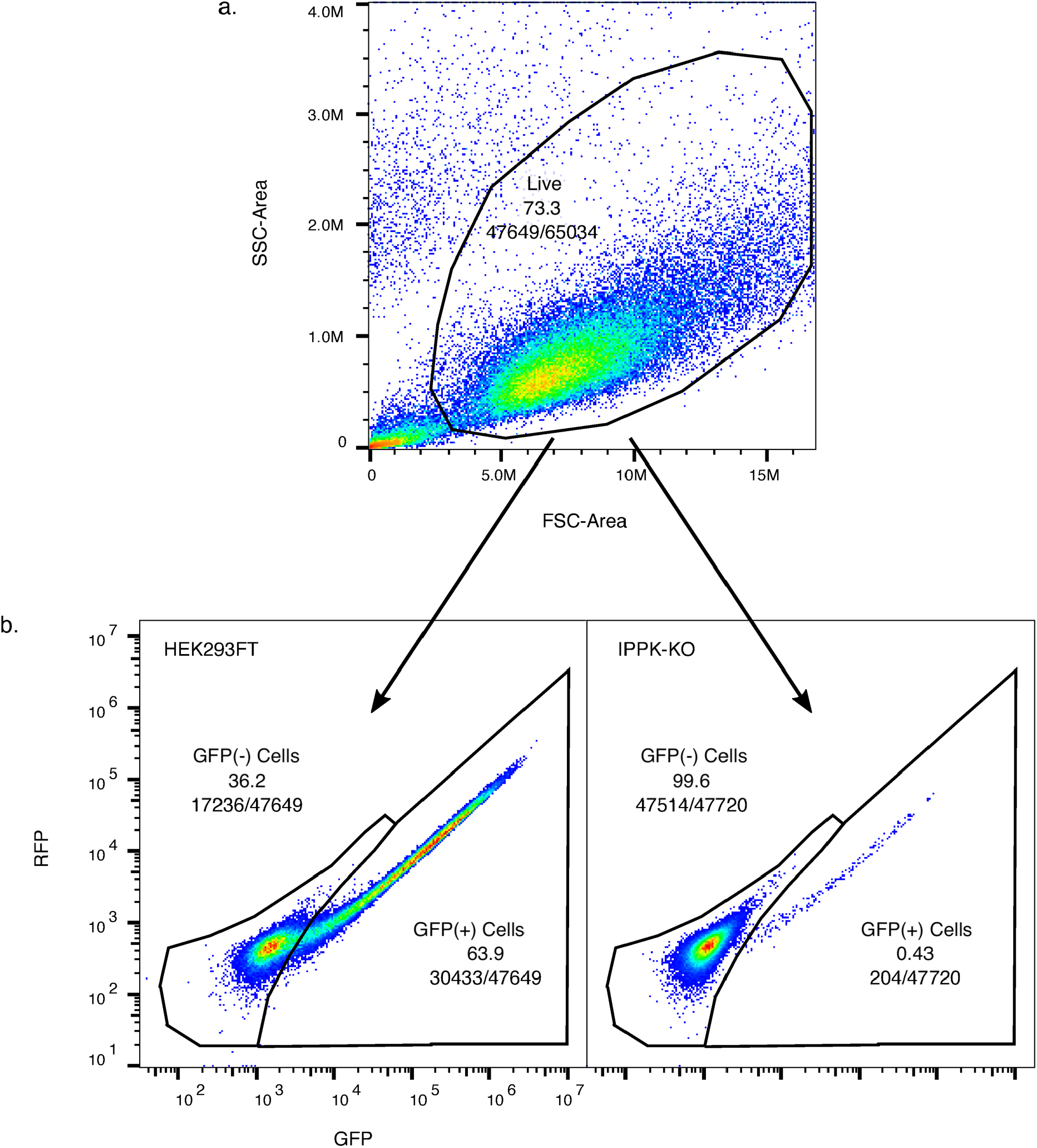
Flow cytometry gating strategy. a) Events were plotted along the forward (FSC) and the side scatter (SSC) x-axis and y-axis respectively. Live cells were gated for the correct size and morphology for further sorting. **b)** Live cells were plotted along GFP and RFP on the x-axis and y-axis respectively. Cells not expressing GFP, GFP(-), and cells expressing GFP, GFP(+), were gated accordingly. Left, a representative plot of virus titer from an RSV-GFP provirus produced in WT HEK293FTs. Right, a representative plot of virus titer from an RSV-GFP provirus produced in IPPK-KO HEK293FTs.

**Figure S10:**
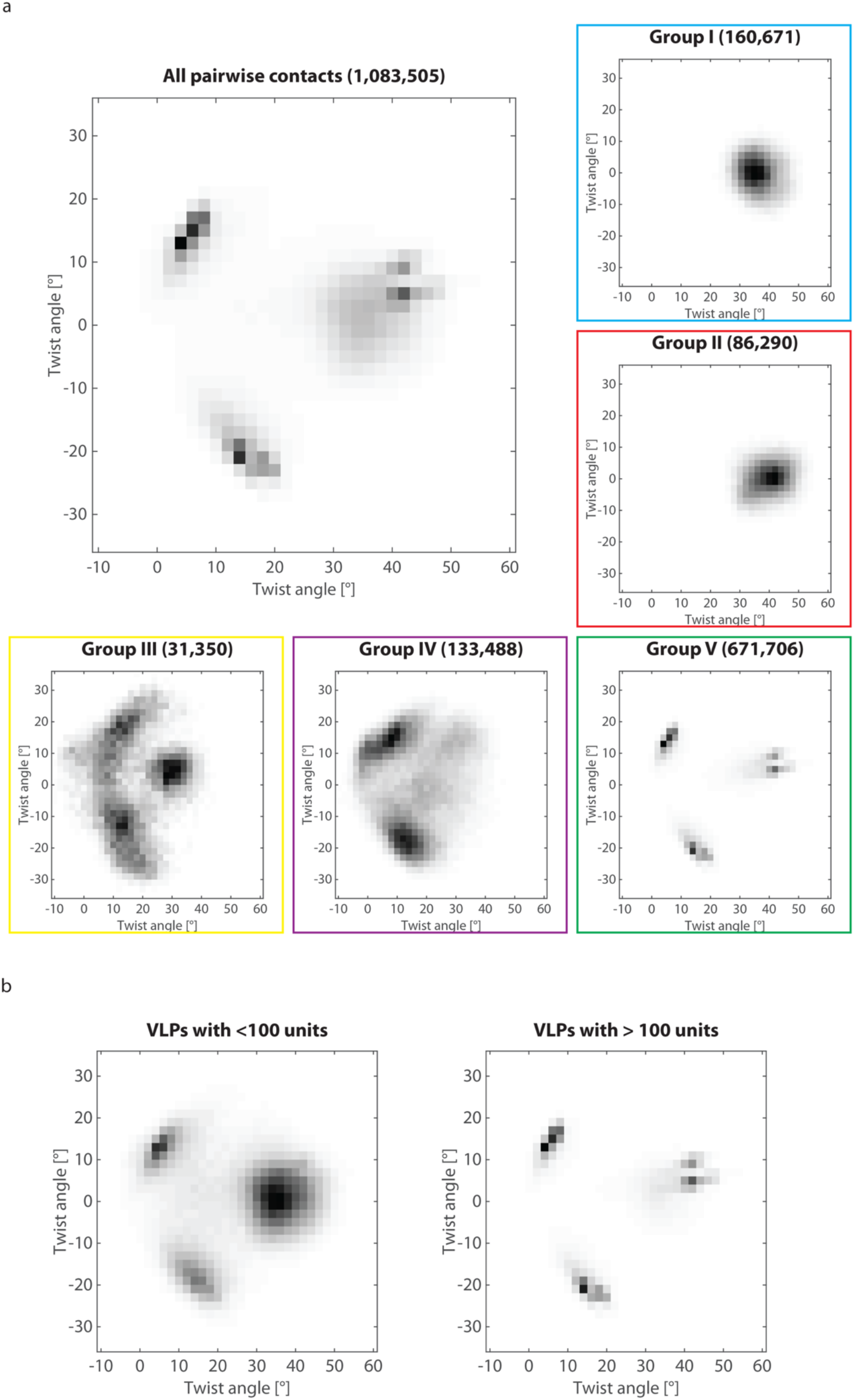
Flexibility of RSV CASPNC VLPs. Tilt-twist plots were used to examine flexibility of unit pairs. Since neither tilt nor twist angles were used as criteria during classification, they could be utilized to assess the quality of classification and progress of subtomogram alignment. **a)** Large panel: all pairwise contacts between adjacent units in the CASPNC dataset; small panels: contacts between unit pairs belonging to the different groups. Coloring is as in Figure 3 and Supplementary Figure 3. **b)** Tilt-twist analysis plots for VLPs with less or more than 100 units are shown on the left and right side, respectively. Note that large VLPs consist mostly of group V – tubular unit connections.

## Supplemental movie legends

**Movie S1: Mature RSV CASPNC tubular and polyhedral VLPs**. Animated Z-slices through an example tomogram. CA hexamers and pentamers in the tomogram are visualized by cyan hexagons and red pentagons, respectively.

**Movie S2: 3D visualization of the structure of RSV CA hexamer derived from tubular VLPs**. Video of the low-pass filtered composite map of an RSV CASPNC tube, the 4.3 Å structure of the RSV CA hexamer, and the refined model (CA_NTD_ – cyan, CA_CTD_ – orange).

**Movie S3: 3D visualization of a composite map of the polyhedral VLP shown in Fig. 3**. The map was built by placing the averages for the respective classes into the positions determined by subtomogram alignment. The color code corresponds to Fig. 3. The structure of the pentamer-hexamer interface solved to 6.0 Å resolution is shown together with the rigid-body fitted model (CA_NTD_ – cyan, CA_CTD_ – orange).

**Movie S4: Morph between hexamer – pentamer, and two hexamers sharing a pentamer neighbor.** Two adjacent CA_NTD_ domains of one hexamer are colored in blue and cyan. Note the amount of movement between the highlighted domains induced by accommodating the pentamer.

**Movie S5: Morph between pentamer-pentamer interface from CA T=1 icosahedron and low-tilt hexamer-hexamer interface from CASPNC tubes.** The color-highlighted interface is stationary compared to the rest of the structure as morphing between the two cryo-EM maps occurs.

## Notes

### Competing Interest Statement

The authors have declared no competing interest.

